# Inhibiting DDX3X triggers tumor-intrinsic type I interferon response and enhances anti-tumor immunity

**DOI:** 10.1101/2020.09.09.289587

**Authors:** Hyeongjwa Choi, Juntae Kwon, Jiafang Sun, Min Soon Cho, Yifan Sun, John L Casey, Jeffrey Toretsky, Cecil Han

## Abstract

Accumulating evidence has shown that cellular double-stranded RNAs (dsRNAs) induce antiviral innate immune responses in human normal and malignant cancer cells. However, it is not fully understood how endogenous ‘self’ dsRNA homeostasis is regulated in the cell. Here, we show that an RNA-binding protein, DEAD-box RNA helicase 3X (DDX3X), prevents the aberrant accumulation of cellular dsRNAs. Loss of DDX3X induces dsRNA sensor-mediated type I interferon signaling and innate immune response in breast cancer cells due to abnormal cytoplasmic accumulation of dsRNAs. Dual depletion of DDX3X and a dsRNA-editing protein, ADAR1 synergistically activates the cytosolic dsRNA pathway in breast cancer cell. Moreover, inhibiting DDX3X enhances the antitumor activity by increasing tumor intrinsic-type I interferon response, antigen presentation, and tumor-infiltration of cytotoxic T cells as well as dendritic cells in breast tumors, which may lead to the development of breast cancer therapy by targeting DDX3X in combination with immune checkpoint blockade.

## INTRODUCTION

Double-stranded RNAs (dsRNAs) within cells are often a result of infected virus replication and considered as a “danger signal” by the host immune system ^1^. Upon recognition of these ‘‘non-self’’ dsRNAs, the host cytoplasmic dsRNA sensors trigger type I Interferon (IFN) production and induce antiviral responses via upregulation of IFN-stimulated genes (ISGs), that eventually leads to cellular apoptosis or growth inhibition ^2,3^. These dsRNA sensors include Toll-like receptor family proteins (TLR3, TLR7), retinoic acid inducible gene I (RIG-I), melanoma differentiation associated gene 5 (MDA5), and protein kinase R (PKR) ^1,4,5^. In addition, another dsRNA binding protein, 2’-5’-oligoadenylate synthetase 1 (OAS1) directly acts on viral RNA by inducing its degradation ^1^.

Advanced transcriptomics studies have shown that metazoan cells express various types of endogenous “self” dsRNAs such as endogenous retroviral elements (ERVs), repetitive RNA elements, mitochondrial dsRNAs, mRNAs with inverted Alu-containing 3’ UTRs, and structural dsRNAs with long dsRNA stem ^6–11^. Current evidence has revealed that abnormal accumulation of the endogenous dsRNAs in mammalian cells could trigger antiviral innate immune response through activation of dsRNA sensing pathways, such as MDA5 or PKR, and that this response could cause chronic inflammation and related human diseases ^3,12,13^. In particular, intracellular dsRNA accumulation in tumors stimulates induction of cancer-derived type I IFN, which enhances anti-tumor immunity ^14–18^. Of note, ERVs constitute more than 8% of the human genome and their bi-directional transcription has been shown to increase the formation of dsRNAs ^19–21^. DNA methylation silences most ERVs in normal somatic cells, but some cancers exhibit loss of ERV DNA methylation and consequent aberrant overexpression of ERVs ^22–25^. In particular, treatment of cancer cells with epigenetic inhibitors increases the expression of ERVs and subsequently induces IFN pathway activation ^14,18,23,26,27^. Despite this strong evidence that endogenous dsRNAs could induce antiviral innate immune responses in human cells, it has been not fully understood how normal and malignant cells prevent cellular ‘self’ dsRNA from being recognized by the cytosolic dsRNA sensors and thereby triggering a potentially detrimental innate immune response.

Human ADARs (adenosine deaminases) are known to be critical regulators of dsRNAs ^7,28^. ADAR1 edits adenosine (A) of the double-stranded regions to inosine (I) (known as A-to-I RNA editing), which results in the disruption of dsRNA structures or the retention of edited dsRNAs in the nucleus ^29,30^. Also, in the mitochondria, RNA degradosome components SUV3 and polynucleotide phosphorylase (PNPase) prevent the accumulation of mitochondrial double-stranded RNAs ^9,10,15^.

DEAD-box RNA helicase 3X (DDX3X) is a member of DEAD-box RNA helicase superfamily 2 ^31,32^. DDX3X utilizes ATP hydrolysis and RNA helicase activities to unwind RNA duplexes, and participates in the multiple aspects of RNA metabolism, such as transcription, RNA splicing, RNA transport, and initiation of translation ^33–36^. DDX3X has been considered as a promising target for anticancer because of its predicted druggability and involvement in tumorigenesis ^37–43^. In particular, increased DDX3X levels have been found in primary and metastatic breast cancer, and this high expression has been correlated with worse survival ^40,44–46^. Nevertheless, the pathological mechanisms by which DDX3X expression decreases cancer survival remain largely unknown.

Here, we reveal that loss of DDX3X leads to the aberrant cytosolic accumulation of endogenous dsRNAs in the breast cancer cells, which triggers intrinsic type I IFN production through the activation of cytoplasmic dsRNA sensing pathway along with resultant anti-proliferative effect. Inhibiting DDX3X expression also enhances the antigen presentation on the cancer cells as well as anti-tumor immunity in the syngeneic breast tumor mouse model. Taken together, these observations suggest that targeting DDX3X could be a novel way of enhancing anti-tumor immunity and thus contribute to combination immunotherapy approaches for patients with breast cancer and other malignancies.

## RESULTS

### Loss of DDX3X increases the expression of genes in antiviral innate immune response in breast cancer cells

High expression of DDX3X mRNA and protein is correlated with worse survival in patients with breast cancer in Kaplan-Meier plot analysis (Extended Data Fig. 1a). To assess the impact of DDX3X on the gene expression in the breast cancer, we stably depleted DDX3X in breast cancer MCF7 cells using a short hairpin RNA (shRNA)-mediated knockdown, and performed a genome-wide transcriptome analysis using a next-generation RNA deep sequencing (Fig. 1a and Extended Data Fig. 1b). Compared to the control MCF7 with shNon-specific RNA (shNS), DDX3X KD (shDDX3X) greatly increased the genes related to antiviral innate immune responses (Fig. 1a and Supplementary Table 1). Ingenuity pathway analysis (IPA) of differentially expressed genes (DEGs) in DDX3X-control versus (vs.) -knockdown (KD) MCF7 cells identified the upregulation of type I interferon signaling, antigen presentation pathway (APP), and pattern recognition receptors (PRRs) response to bacteria and virus (Fig. 1b). In the gene-set enrichment analysis (GSEA) of DEGs, the top 5 gene sets co-enriched in the DDX3X KD cells included type I interferon (IFN), antigen presentation via major histocompatibility complex (MHC) class I, and viral defense response (Fig. 1c).

**Fig. 1:**
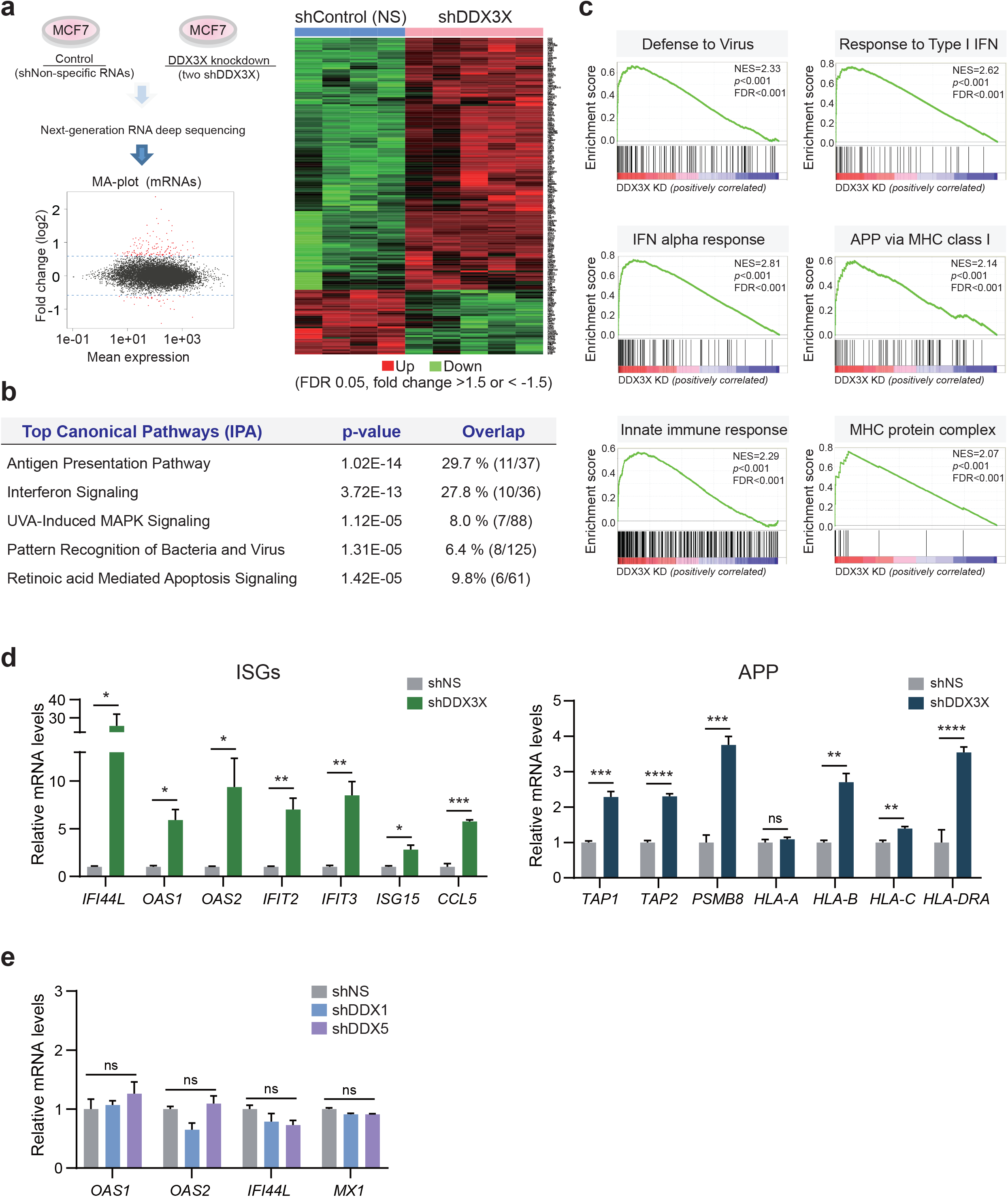
Loss of DDX3X increases the expression of genes in antiviral innate immune response in breast cancer cells. **a**, Genome-wide transcriptome analysis in DDX3X-control (shNS) or -knockdown (shDDX3X) MCF7 cells using a next generation RNA deep sequencing. DEGs (FDR 0.05, fold change >1.5 or < −1.5) in shNS vs. shDDX3X are shown in a MA-plot (labeled in red) and in a heat map. **b**, IPA analysis shows top canonical pathways of DEGs in DDX3X-control or -knockdown (KD) MCF7. **c**, GSEA analysis of DEGs in DDX3X-control or -KD MCF7. **d**, qRT-PCR of mRNA expression of ISGs and APP in DDX3X-control or -KD MCF7 cells. **e**, qRT-PCR of ISGs expression in DDX1-KD or DDX5-KD MCF7 cells, respectively. Data in D and E are representative of three independent experiments and presented as mean ± SEM. Statistics were calculated using unpaired t-tests. \**P* < 0.05; \*\**P* < 0.01; \*\*\**P* < 0.001; ns, not significant.

Using qRT-PCR analysis, we validated that DDX3X KD increased the gene expression of interferon-stimulated genes (ISGs: *IFI44L*, *OAS1*, *OAS2*, *IFIT2*, *IFIT3*, *ISG15*) and T cell chemoattractant (*CCL5*) (Fig. 1d). Also, DDX3X KD upregulated the genes related to antigen processing and presentation (APP) such as MHC class molecules (*HLA-B*, *HLA-C*, and *HLA-DRA*), peptide transporters (*TAP1* and *TAP2*), and immunoproteasome (PSMB8) (Fig. 1d). We observed the similar upregulation of ISGs in the siRNAs-mediated transient DDX3X KD in MCF7 cells (Extended Data Fig. 1c). Knockdown of other DEAD-box helicases such as DDX1 or DDX5 did not present a similar induction of ISGs, suggesting the specific role of DDX3X in the regulation of innate immune responses (Fig. 1e and Extended Data Fig. 1d). Similar upregulation of ISGs was validated in DDX3X-depleted breast cancer MDA-MB-453 cells (Extended Data Fig. 1e). This increased ISGs expression suggests the global upregulation of an IFN-driven transcriptional activation in the DDX3X-depleted breast cancer cells.

### Type I IFN production and STAT1 pathway are activated in DDX3X-depleted cancer cells

To determine if DDX3X knockdown (KD) induces IFN production, we measured the levels of the type I IFN (interferon-alpha, α; interferon-beta, β) and type II IFN (interferon-gamma, γ) in the culture media of the DDX3X KD cancer cells. DDX3X KD significantly induced the expression and secretion of the type I IFN-α and IFN-β, but not type II IFN-γ in the human and mouse breast cancer cells, including MCF7, MDA-MB-453, and E0771 (Fig. 2a). We also observed the increased production of IFN-α and IFN-β in the DDX3X-depleted melanoma cell A375 (Extended Data Fig. 2a).

**Fig. 2:**
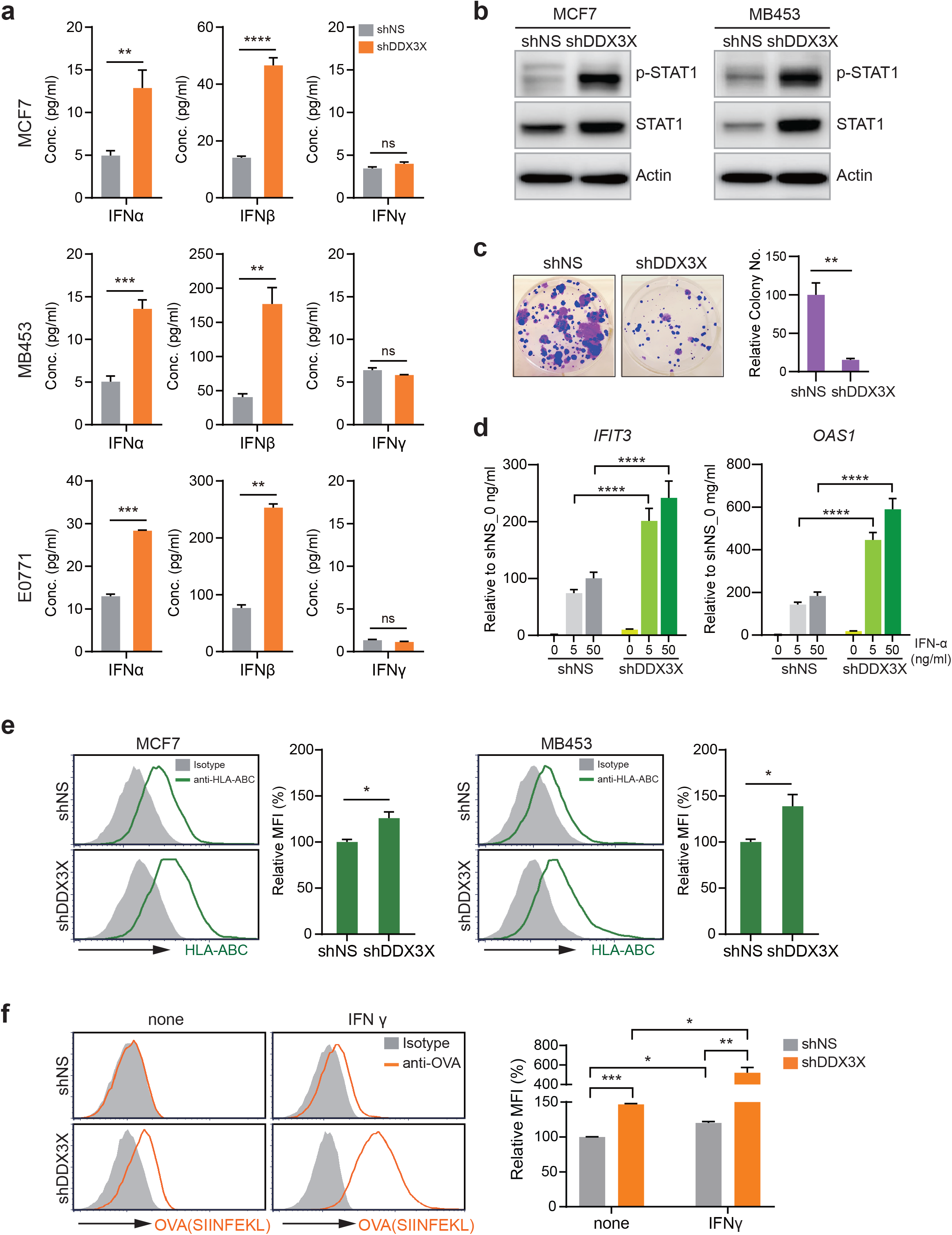
Loss of DDX3X activates type I IFN production, suppresses cell proliferation, and induces antigen presentation in cancer cells. **a**, ELISA of IFN-α, -β, and -γ in the culture supernatants from DDX3X-control or -depleted MCF7, MB453, and E0771 cells, respectively. **b**, Western blot analysis of STAT1 and phosphorylated STAT1 in the DDX3X-control or -depleted MCF7 and MDA-MB-453 cells, respectively. **c**, Colony formation assay in DDX3X-control or -KD MCF7 cells shows cell growth inhibition by DDX3X depletion. **d**, qRT-PCR of *IFIT3* and *OAS1* in DDX3X-control or -KD MCF7 cells after IFN-α treatment for 5 hours. **e**, Representative flow histograms (left) and a relative mean fluorescence intensity (MFI) quantification bar graph (right) of HLA-ABC expression in the DDX3X-KD MCF7 or DDX3X-KD MDA-MB-453 cells, respectively. **f**, Representative histograms (left) and a relative MFI quantification bar graph (right) of SIINFEKL bound to H-2Kb expression on DDX3X deficient B16-OVA cells. Cells were treated 100 ng/ml of IFNγ for 48 hours. Data are representative of three independent experiments. Data are shown as mean ± SEM of three independent experiments. Statistics were calculated using unpaired t-tests. \**P* < 0.05; \*\**P* < 0.01; \*\*\**P* < 0.001; \*\*\*\**P* < 0.0001; ns, not significant.

Canonical type I IFN signaling leads to the transcription of hundreds of IFN-stimulated genes (ISGs) through the activation of Janus kinase-signal transducer and activator of transcription (JAK/STAT) pathway ^2^. To determine whether the loss of DDX3X could activate the JAK/STAT pathway, we assessed the components of the STAT pathway. We confirmed that MCF7 and MDA-MB-453 express the IFN receptors and have an intact JAK/STAT pathway in response to type I IFN treatment (Extended Data Fig. 2b,c). DDX3X KD increased the level of the phosphorylated STAT1 and total STAT1 (Fig. 2b). The phosphorylated STAT2 was not detected, but DDX3X KD increased total STAT2, which belongs to ISGs (Extended Data Fig. 2d). The activated antiviral response often results in the induction of the apoptosis or growth inhibition in the cells ^2^. DDX3X KD did not induce apoptosis (Extended Data Fig. 2e), but significantly suppressed the cell proliferation (Fig. 2c and Extended Data Fig. 2f). Taken together, these data suggest that the depletion of DDX3X in breast cancer cells significantly induces cancer cell production of type I IFNs that drives ISG expression through the STAT1 pathway, and leads to an anti-proliferative effect.

### Loss of DDX3X enhances antigen presentation and sensitizes breast cancer cells to IFN treatment

Next, we observed that IFN-α treatment more dramatically increases the ISG expression in the DDX3X KD than control MCF7 cells, showing that inhibiting DDX3X sensitizes the cancer cells to type I IFN treatment (Fig. 2d). To further determine the functional consequences of the increased IFN-α and IFN-β production on the antigen presentation of DDX3X KD cells, we assessed the cell surface expression of MHC class I proteins using flow cytometry. DDX3X-depleted MCF7 cells displayed the significantly enhanced expression of MHC class I proteins (HLA-ABC) (Fig. 2e). Although qRT-PCR analysis showed the increased gene expression of *HLA-DRA*, one of the subunits of MHC class II molecule (Fig. 1d), we did not see the increased MHC II protein complex on the DDX3X KD cells (Extended Data Fig. 2g). We also stably knocked-down DDX3X in the ovalbumin (OVA)-expressing mouse cancer cells, B16-OVA. DDX3X KD increased the presentation of the OVA epitope (SIINFEKL) derived from OVA in MHC I molecules on the cell surface (Fig. 2f). The OVA peptide presentation was dramatically increased in the DDX3X KD cells by IFN-γ stimulation (Fig. 2f). This data indicates that DDX3X KD enhances the antigen processing as well as antigen presentation via MHC I on the cancer cells.

### Inhibiting DDX3X triggers the accumulation of endogenous double-stranded RNAs

Failure to degrade the excess endogenous double-stranded RNAs (dsRNAs) is known to activate the cytoplasmic dsRNA sensors and trigger type I IFN response in the several types of cancer cells ^3,7,9,12^. Because DDX3X is an RNA-binding protein, which has an activity for dsRNA unwinding, we speculated that DDX3X could impact the level of endogenous dsRNAs. To measure endogenous dsRNA in the cells, we used a monoclonal anti-dsRNA specific J2 antibody (Scion, Hungary), which is widely used to recognize viral and cellular dsRNA of more than 40-bp length with no sequence specificity in animals and plants ^47–49^.

First, we validated the suitability of the J2 antibody to detect endogenous dsRNA in our systems, including immunostaining, dot blot, northwestern blot, and flow cytometry. A dsRNA-specific J2 antibody specifically detected the dsRNA signals in the various cancer cells. The J2 signals were sensitive to the dsRNA-specific RNase III or RNase A, but not affected by the treatment of the single-stranded RNA (ssRNA)-specific RNase I or DNase I (Extended Data Fig. 3a, b). J2 antibody also recognized the synthetic dsRNA mimic analog, polyinosinic–polycytidylic acid (poly (I:C)) *in vitro* and in the poly I:C transfected cells (Extended Data Fig. 3c). DNA methyltransferase inhibitors are known to increase the bi-directional transcription of endogenous retroviral elements (ERVs) or repetitive RNA elements, which could increase the RNA duplexes (Extended Data Fig. 3d) ^18,26,50^. J2 antibody was able to distinguish the increased level of endogenous dsRNAs in response to the treatment of DNA methyltransferase inhibitor, 5-AzaC in the several cancer cells using dot blot and flow cytometry approaches (Extended Data Fig. 3e, f). Collectively, these data validate that the J2 antibody specifically recognizes the authentic cellular dsRNAs in cancer cells.

Next, using the validated anti-dsRNA specific J2 antibody, we investigated the level of endogenous dsRNAs in the DDX3X-depleted breast cancer cells. We found that depletion of DDX3X substantially increased the level of endogenous dsRNAs in MCF7 cells using immunostaining (Fig. 3a), flow cytometry (Fig. 3b), northwestern blot (Fig. 3c), and dot blot analysis approaches (Fig. 3d and Extended Data Fig. 3g). Enhanced dsRNA signals were also detected in the human and mouse DDX3X KD cancer cells (MDA-MB-453, A375, 4T1, B16F10) (Fig. 3e). Next, We checked the transcription level of the ERVs that were previously identified ^18,25,26,50^. DDX3X-depleted MCF7 cells showed the increased expression of several ERVs (Fig. 3f). Using TAG-aided sense/antisense transcript detection PCR (TASA-TD PCR) that detects the bi-directional expression of ERVs, we validated the dsRNA form of ERVs (Fig. 3g). Similarly, increased ERV expression was detected in the DDX3X-depleted MDA-MB-453 cells (Fig. 3h). HEK293T cells are known to be defective in their ability to produce type I IFN in response to viral RNAs or poly I:C stimulation ^51^. There was no induction of ISGs or IFN-β, but dsRNAs still accumulated in the DDX3X KD HEK293T cells (Extended Data Fig. 3h, i), indicating that dsRNA accumulation is a direct consequence of DDX3X loss, rather than IFN activation. In addition to the J2 antibody, we also confirmed similar results using another dsRNA-specific K1 antibody (Scion, Hungary) ^48^ (data not shown). Taken together, these data indicate that DDX3X prevents dsRNA accumulation in the human and mouse cells, suggesting the functional association of DDX3X in regulating the level of endogenous dsRNAs.

**Fig. 3:**
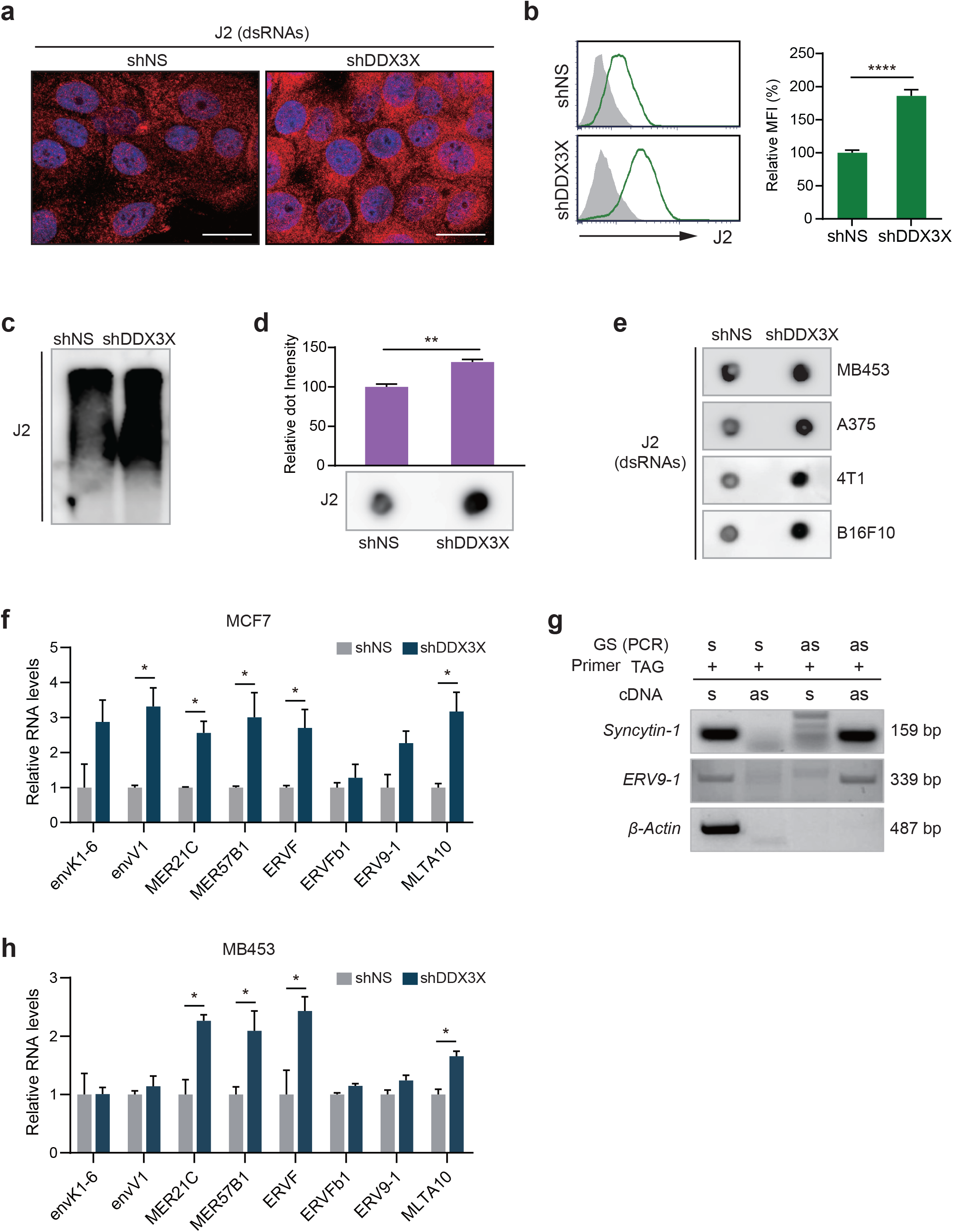
DDX3X inhibition triggers endogenous dsRNAs accumulation. **a**–**d**, Endogenous dsRNA accumulation in DDX3X-control or -KD MCF7 cells analyzed by immunofluorescence (**a**), flow cytometry (**b**), northwestern blot (**c**), and dot blot (**d**) with anti-dsRNA specific (J2) antibody. Scale bars, 25 *μm*. **e**, dsRNA accumulation in DDX3X-depleted MDA-MB-453, A375, 4T1, and B16F10 cells assessed by dot blot. **f**, qRT-PCR of ERV genes in DDX3X-depleted MCF7 cells. **g**, TASA-TD PCR amplified sense and antisense transcripts of the *Syncytin-1* and *ERV9-1* genes. A *β-Actin* used for a sense transcript amplification. *Syncytin-1 used for a positive control for a bi-directional transcript.* **h**, qRT-PCR of ERV genes in DDX3X-control or -KD MDA-MB-453 cells. Data are representative of three independent experiments. Data are shown as mean ± SEM. Statistics were calculated using unpaired t-tests. \**P* < 0.05; \*\**P* < 0.01; \*\*\*\**P* < 0.0001.

### Inhibiting DDX3X activates cytoplasmic dsRNA response signaling in cancer cells

Next, we investigated which signaling pathway is responsible for type I IFN production in the DDX3X-KD breast cancer cells. Pattern recognition receptors (PPRs) recognize the intracellular double-stranded DNAs (dsDNAs) or dsRNAs, and subsequently induce type I IFN production ^1^. First, we examined cytosolic DNA sensing-mediated innate immune response through STING pathway ^52^. There was no activation in cyclic GMP-AMP synthetase (cGAS), a central receptor of cytosolic DNA, and its downstream target, STING (Fig. 4a). This indicates no involvement of a cytosolic DNA, and that DDX3X KD induces type I IFN in a STING-independent manner.

**Fig. 4:**
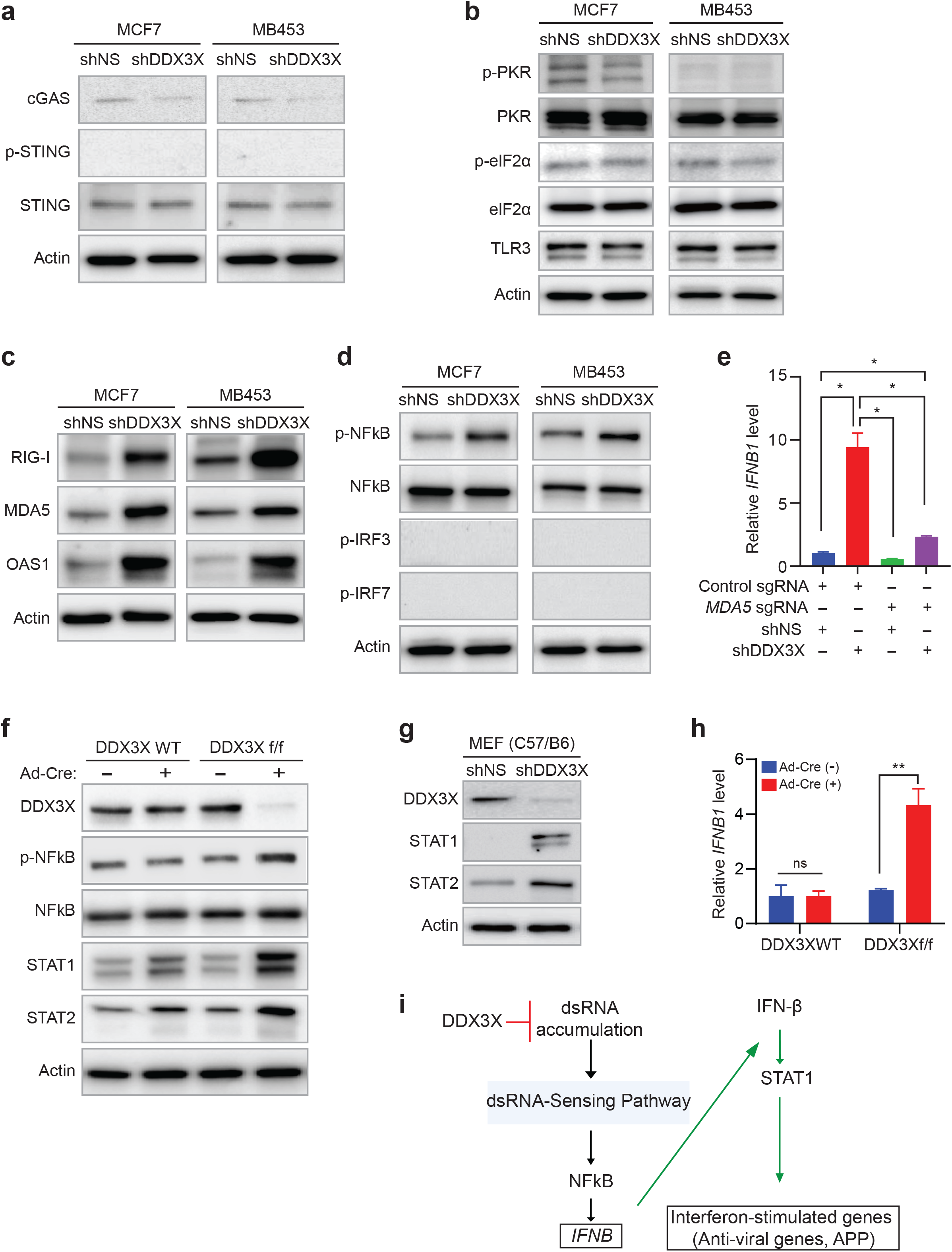
Inhibiting DDX3X activates cytoplasmic dsRNA response signaling in cancer and MEF cells. **a**, Western blot analysis of dsDNA sensor and downstream target, cGAS and STING, in DDX3X-control or -depleted MCF7 or MDA-MB-453 cells. **b**, Western blot analysis of dsRNA sensors and downstream target, TLR3, PKR and elF2a, in DDX3X-control or -depleted MCF7 or -MDA-MB-453 cells. **c**, Western blot analysis of dsRNA sensors, RIG-I, MDA5, and OAS1, in DDX3X-control or - depleted MCF7 or MDA-MB-453 cells. **d**, Western blot analysis of phosphorylation of transcription factors, NFκB, IRF3, and IRF7, in DDX3X-control or -depleted MCF7 or MDA-MB-453 cells. **e**, qRT-PCR of *IFNB1* in DDX3X-control or -KD MCF7 following by MDA5 knockout. **f**, Western blot analysis of phosphorylated and total NFκB, STAT1, and STAT2 in DDX3X wildtype and DDX3X^f/f^ MEF cells after Ad-Cre virus treatment. **g**, Western blot analysis of STAT1 and STAT2 in shRNA-mediated DDX3X KD MEF cells. **h**, qRT-PCR of *IFNB1* in DDX3X^f/f^ MEF cells 4 days after Ad-Cre virus treatment. **i**, Schema of the dsRNA-mediated type I IFN signaling in DDX3X-depleted cancer and MEF cells. Data are representative of three independent experiments. Data are shown as mean ± SEM. Statistics were calculated using unpaired t-tests. \**P* < 0.05; \*\**P* < 0.01; ns, not significant.

Because the loss of DDX3X resulted in the accumulation of dsRNAs (Fig.3), we investigated dsRNA-sensing pathways. First, we examined several intracellular dsRNA sensors, including PKR, TLR3, OAS1, RIG-I (DDX58), and MDA5 (IFIH1). The levels of TLR3, phosphorylated PKR, and phosphorylated eIF2α were not changed, indicating that there was no PKR or TLR3 activation by DDX3X depletion (Fig. 4b). However, the protein and mRNA expression of MDA5, RIG-I, and OAS1 were significantly increased in the DDX3X KD cells (Fig. 4c and Extended Data Fig. 4a, b). Next, we investigated if there were changes in the transcription factors (IRF3, IRF7, NFκB) that are responsible for IFN-β production in DDX3X KD cells. Phosphorylated NFκB was consistently increased in the DDX3X-depleted cancer cells (Fig. 4d). DDX3X KD MCF7 cells showed no phosphorylation of IRF3 or IRF7 (Fig. 4d), although MCF7 cells have an intact activation of IRF3 and IRF7 in response to poly I:C treatment (Extended Data Fig. 4c). CRISPR/Cas9-mediated MDA5 knockout significantly diminished the induction of IFN-β expression by the DDX3X depletion in the MCF7 cells (Fig. 4e and Extended Data Fig. 4d). This suggests that MDA5 is the critical dsRNA sensor to initiate the dsRNA-mediated type I IFN production in the DDX3X-depleted MCF7 cells. Collectively, these data imply that inhibiting DDX3X increases the endogenous dsRNA abundance, which activates the cytosolic dsRNA sensing pathway, mainly the MDA5-NFκB signaling axis, that leads to type I IFN production in breast cancer cells.

### Type I IFN and dsRNAs signaling pathway are also activated in the DDX3X-null MEF cells

Furthermore, we investigated if the loss of DDX3X could activate dsRNA signaling in mouse embryonic fibroblast (MEF) cells. We generated a *Ddx3x* conditional knockout mouse (named DDX3X^f/f^), in which Ddx3x exons 5~7 were floxed by two loxP sites (Extended Data Fig. 4e, f). MEF cells were isolated from E12.5 embryos by breeding DDX3X^f/+^ (or DDX3X^f/f^) and DDX3X^Y/f^ mice, and treated with Cre recombinase expressing adenovirus (Ad-Cre) to induce Cre-mediated *Ddx3x* gene deletion. Both hemizygous and homozygous DDX3X-null MEF cells (*Ddx3x*^Y/f^ male and *Ddx3x*^f/f^ female) showed the increased total STAT1 and STAT2 levels (Extended Data Fig. 4g). Heterozygous DDX3X knockout (*Ddx3x*^f/+^ female) MEF cells showed the less significant increased STAT1 and no induction of STAT2 level after Ad-Cre treatment (Fig. 4f and Extended Data Fig. 4g). Consistent with the Cre-mediated DDX3X-null MEF cell, shRNA-derived DDX3X knockdown MEF cells also showed the increased level of STAT1 and STAT2, while the non-specific shRNA did not affect the level of STAT1 and STAT2 (Fig. 4g). Consistent with the cancer cells, the loss of DDX3X in the MEF cells also showed the increased phosphorylation of NFκB (Fig. 4f). Importantly, *IFN-β* mRNA expression was increased in the DDX3X-null MEF cells (Fig.4h and Extended Data Fig. 4h). Taken together, these data suggest that DDX3X prevents the activation of cellular dsRNA signaling and type I IFN response in the human cancer cells and mouse embryonic fibroblast (Fig. 4i).

### DDX3X interacts with cytoplasmic ADAR1 and prevents the buildup of cytoplasmic dsRNA

To understand how DDX3X depletion could activate the cytosolic dsRNA sensors, we first assessed the nuclear-cytoplasmic distribution of dsRNAs in DDX3X-control or -KD MCF7 cells. We isolated total RNAs from the nuclear and cytoplasmic fractions of the DDX3X-control or -KD MCF7 cells, respectively, and applied the isolated RNAs to the dot blot analysis with a dsRNA-specific J2 antibody (Fig. 5a and Extended Data Fig. 5a). Most of dsRNA was found in the nuclear fraction, despite the fact that the amount of RNAs used for dot blot analysis was much higher in the cytoplasmic fraction than the nuclear fraction (Fig. 5a and Extended Data Fig. 5b). DDX3X KD not only increased the level of dsRNA in the nuclear fraction but also substantially increased the cytoplasmic level of dsRNAs (4.3% to 19.3%) (Fig. 5a and Extended Data Fig. 5b). This suggests that DDX3X regulates dsRNAs in both nucleus and cytoplasm. Next, we determined whether this increased cytosolic dsRNAs are the result of mitochondrial dsRNA (mtdsRNA) leaking from the mitochondria. We did not see any increased mitochondrial RNAs in the cytosolic fraction of DDX3X KD cells compared to the control cells, suggesting the accumulated endogenous dsRNAs mainly originated from the genomic DNA, not from the mitochondria (Extended Data Fig. 5c).

**Fig. 5:**
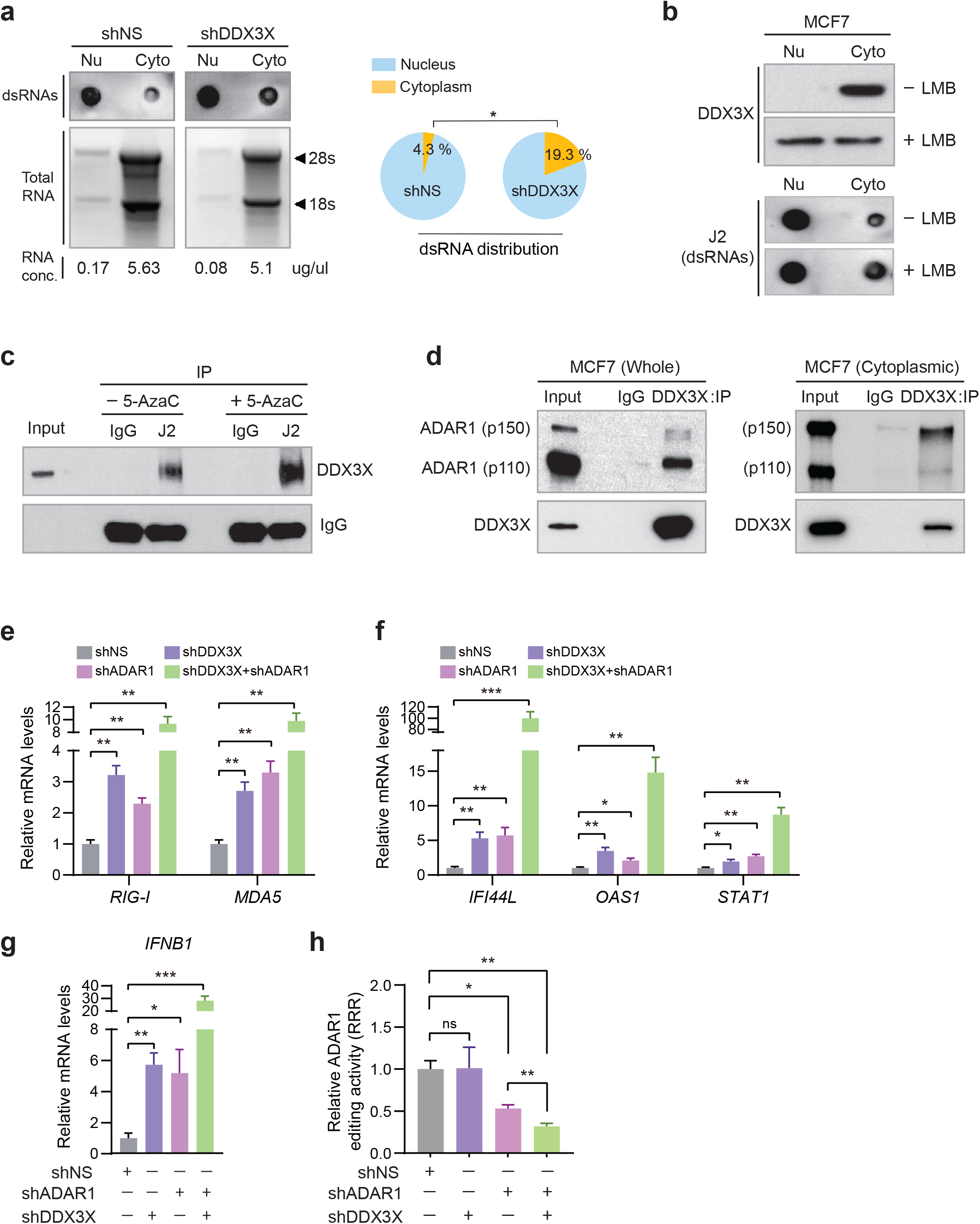
DDX3X interacts with cytoplasmic ADAR1 and prevents the buildup of the cytoplasmic dsRNA. **a**, J2 dot blot analysis of RNA extracts from cytoplasmic or nucleus fractions of the DDX3X-control or -KD MCF7 cells. Electrophoresis analysis of RNAs isolated from cytoplasmic or nucleus fractions. A circle graph shows relative dot intensity (each cells’ cytoplasm + nucleus = 100%). **b**, Localization of DDX3X (upper) and levels of dsRNAs (lower) in cytoplasmic or nucleus fractions after leptomycin B (LMB; 30nM, 16 h) treatment. **c**, J2 Immunoprecipitation with whole MCF7 cell extracts with or without 5-AzaC treatment. MCF7 cells were fixed with 1 % formaldehyde and IP was performed. dsRNA-bound proteins were analyzed using western blot with anti-DDX3X or anti-IgG (light chain) antibodies. **d**, Interaction of DDX3X and ADAR. Immunoprecipitation of DDX3X from whole cell extracts or cytoplasmic fraction of MCF7 cells. IP was performed with DDX3X antibody or control IgG, respectively. Western blot analyzed using anti-ADAR1 and anti-DDX3X antibodies. **e**–**g**, mRNA expression of dsRNA sensing genes (**e**), ISGs (**f**), and *IFNB1* (**g**) in DDX3X and ADAR1 single, or double KD cells. **h**, Determination of ADAR1 editing as the ratio between luminescence from Nluc/FFL expressed reporter plasmid. Data was calculated a relative response ratio (RRR) = (experimental sample ratio-negative control ratio) / (positive control ratio-negative control ratio)). Data are representative of three independent experiments. Data represented as mean ± SEM. Statistics were calculated using unpaired t-tests. \**P* < 0.05; \*\**P* < 0.01; \*\*\**P* < 0.001.

DDX3X is known to be a shuttling protein between the cytoplasm and nucleus whose steady-state localization is mostly cytoplasmic ^53,54^, When CRM1-dependent nuclear export was blocked with the drug leptomycin B ^55^, DDX3X was stuck in the nucleus, and subsequently, the cytoplasmic level of DDX3X (60% reduction) was decreased (Fig. 5b). Interestingly, when we blocked the nuclear export with leptomycin B, the level of cytoplasmic dsRNAs was increased in the cytoplasm while the nucleus dsRNA level was unchanged (Fig. 5b). This data suggests that the level of cytoplasmic DDX3X is important to maintain the endogenous dsRNAs in the cytoplasm at a minimum level. Next, to further determine whether DDX3X is directly involved in regulating the level or stability of endogenous dsRNAs in cells, we examined if DDX3X is directly associated with endogenous dsRNAs in the cells. We carried out immunoprecipitation to pull-down dsRNA with a dsRNA-specific J2 antibody in the MCF7 cells after treating with formaldehyde to generate cross-links between RNA and protein. DDX3X was co-precipitated with dsRNAs, but not with the control IgG (Fig. 5c). Furthermore, 5-AzaC treatment, which induces the transcription of endogenous dsRNAs, enhanced the binding of DDX3X with dsRNAs (Fig. 5c). This indicates that DDX3X is directly associated with endogenous dsRNAs in the cells.

In particular, DDX3X specifically interacts with ADAR1’s two isoforms (110 kDa, ADAR1-p110; 150 kDa, ADAR1-p150) in MCF7 cells (Fig. 5d and Extended Data Fig. 5d). ADAR1 edits dsRNA (from Adenosine to Inosine) mostly in the nucleus, and most ADAR1-p110 is found in the nucleus ^7,56^. Because DDX3X has been found mainly in the cytoplasm (Extended Data Fig. 5e), we further dissected the subcellular localization for the interaction between DDX3X and ADAR1. We found that DDX3X interacted with primarily ADAR1-p150 in the cytoplasm (Fig. 5d, right panel). Notably, DDX3X KD alone increased the levels of dsRNA sensors (*RIG-I*, *MDA5*), ISGs (*IFI44L*, *OAS1*, *STAT1*), and IFN beta comparable to the effect of ADAR1 KD alone (Fig. 5e-g and Extended Data Fig. 5g). Moreover, the double knockdown of ADAR and DDX3X greatly increased the expression of *RIG-I*, *MDA5*, other ISGs as well as *IFN beta* (Fig. 5e-g). To further explore if DDX3X directly impacts on ADAR1-mediated A-to-I dsRNA editing, we used a dual-luciferase reporter for monitoring ADAR-editing efficiency (gift from Dr. Ohman) ^57^. ADAR1-mediated dsRNA editing activity was determined as the ratio between luminescence from the Nano luciferase/Firefly luciferase (Extended Data Fig. 5h). We transiently expressed a negative (0% editing), a positive (100% editing), and an editing reporter (A can be edited to I) vectors, respectively, to validate ADAR1 editing activity in the different parental cell lines (Extended Data Fig. 5i). We found that DDX3X KD alone in MCF7 cells did not impact on the activity of ADAR1-mediated A-to-I editing (Fig. 5h). However, the double knockdown of ADAR1 and DDX3X significantly reduced the dsRNA editing activity relative to ADAR1 KD alone (Fig. 5h). These data indicate that double-inhibition of DDX3X and ADAR1 synergistically activates the cytosolic dsRNA-signaling pathway and an innate immune response in the cancer cells.

### Low DDX3X level is related to the chronic activation of the innate antiviral immune response in the various cancer cells

To explore DDX3X-dsRNAs-type I IFN signaling axis in other solid cancers, we compared the gene expression profiles in the DDX3X-highly expressing cancer cells (DDX3X^hi^, 253 cell lines) vs. DDX3X-low expressing cells (DDX3X^low^, 253 cell lines) of the Cancer Cell Line Encyclopedia (CCLE) database (Extended Data Fig. 6a). The DDX3X expression in both DDX3X^hi^ and DDX3X^low^ groups was not affected by sex or origin of the tumor types (Extended Data Fig. 6b, c). We identified 386 differentially expressed genes (DEGs) in DDX3X^hi^ vs. DDX3X^low^ cancer cells (cut off at *p*-value < 0.01 and fold change > 1.5). Forty-nine genes in the DDX3X^hi^ cell group and 337 genes in the DDX3X^low^ cell group were upregulated, respectively (Fig. 6a).

**Fig. 6:**
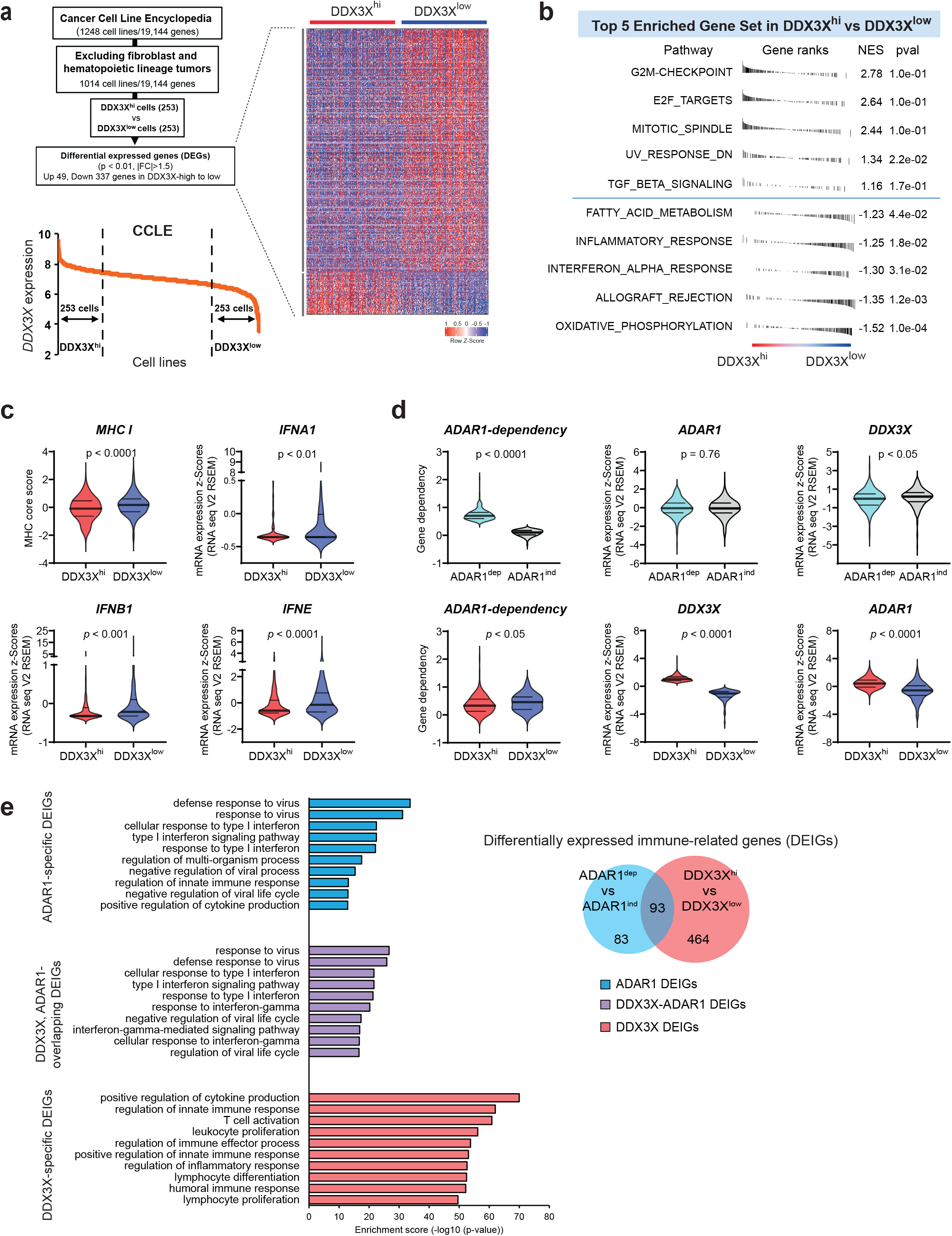
Low DDX3X level is related to the chronic activation of the innate antiviral immune response in the various cancer cells. **a**, Flowchart detailing the steps leading to the identification of differentially expressed genes (DEGs) between DDX3X^hi^ and DDX3X^low^ cancer cells in CCLE (left). DDX3X^hi^ (top 25%) and DDX3X^low^ (bottom 25%) cell line groups based on DDX3X expression. Heatmap of 386 DEGs in DDX3X^hi^ vs. DDX3X^low^ cancer cells (right). **b**, Gene set enrichment analysis (GSEA) plots for 5 top scoring hallmark gene set collection from the Molecular Signatures Database (MSigDB). **c**, Violin plots showing MHC I and type I IFN transcript levels in DDX3X^hi^ and DDX3X^low^ group. **d**, Comparison of gene dependency and expression level of ADAR1 and DDX3X in the ADA1R^dep^ vs. ADAR1^idp^ (top panel) and DDX3X^hi^ vs. DDX3X^low^ (bottom panel). Gene dependency score is obtained from DeMap portal (Genetic dependency by combined RNAi). A higher score indicates that a gene is more likely to be dependent for survival. **e**, Enrichment of biological processes (GO) terms in each group of differentially expressed immune-related genes (DEIGs) between ADA1R^dep^ vs. ADAR1^idp^ and DDX3X^hi^ vs. DDX3X^low^ groups (FDR<0.05). Bars indicate statistical significance shown as −log10 of *p* value. Venn diagram depicting the number of DEIGs between two groups (right).

In the top 5 enriched gene sets identified in DEGs between DDX3X^hi^ vs. DDX3X^low^, the DDX3X^low^ cell group revealed the significant gene enrichment in the innate immune responses, such as inflammatory response, interferon-α response, and allograft rejection (Fig. 6b). On the other hand, UV response, E2F target, mitotic spindle, and G2M checkpoint were enriched in the DDX3X^hi^ group (Fig. 6b). In line with the finding that DDX3X inhibition induces the type I IFN and MHC class I expression in the breast cancer cells, the elevated level of MHC class genes, types I IFN (*IFNA1*, *IFNB1*, *IFNK*, *IFNE*), and proinflammatory cytokines (*IL6*, *IL1*) were found in the DDX3X^low^ cells (Fig. 6c and Extended Data Fig. 6d, e). This CCLE data suggests that DDX3X expression level is inversely correlated with the increased intrinsic type I IFN and innate immune responses in the various cancer types.

We further investigated the functional relationship between DDX3X and ADAR in the endogenous dsRNA-mediated innate immune response in various cancer cells. Previous studies reported that ADAR1 knockdown kills certain cancer cells, and that STING dependently elevated-intrinsic Type I IFN signaling confers a dependency of these cancer cells on ADAR1 ^58,59^. We selected ADAR1-dependent (ADAR1^dep^, 207 cell lines) and -independent cancer cell (ADAR1^ind^, 207 cell lines) groups from the DRIVE dataset ^60,61^ (Extended Data Fig. 6a). As expected, the ADAR1^dep^ cell group has a high dependency score for the *ADAR1* gene (Fig. 6d, top left panel) (score 0 means a minimal impact of knockdown or knockout of a gene for cell survival; score close to 1 means an essential gene for cell survival). ADAR1^dep^ cell group also showed elevated expression of MHC I and type I IFN (Extended Data Fig. 6f). Interestingly, the ADAR1 level was not different in between ADAR1^dep^ and ADAR1^ind^ cells, indicating the transcript level of ADAR1 does not impact on the ADAR1 dependency (Fig. 6d, top middle panel). ADAR1^dep^ cells showed a lower level of DDX3X than the ADAR1^ind^ cells (Fig. 6d, top right panel). DDX3X^low^ group showed a higher ADAR1 dependency than the DDX3X^hi^ group (Fig. 6d, bottom left panel).

Next, we compared the differentially expressed immune-related gene (DEIG) profiles in the ADAR1-specific DEIG (ADAR1^dep^ vs. ADAR1^ind^) and DDX3X-specific DEIG (DDX3X^hi^ vs. DDX3X^low^). We performed GO pathway analysis of ADAR1-specific DEIG, DDX3X-specific DEIG, and co-identified DEIGs in both DDX3X and ADAR1, respectively (Fig. 6e and Extended Data Fig. 6a). Among the top 10 enriched GO pathways, overlapping DEIGs from the DDX3X-specific and ADAR1-specific DEIG genes (indicated as a purple color) demonstrated the enrichment of innate immunity, including defense response to virus, type I interferon signaling pathway, and negative regulation of viral life cycle (Fig. 6e). Especially, DDX3X-specific DEIGs (indicated as a red color) are additionally related to genes involved with T cell activation, leukocyte proliferation, and positive regulation of cytokine production (Fig. 6e). We observed similar results in GO pathway analysis with the total DEGs from ADAR1^dep^ vs. ADAR1^ind^ and DDX3X^hi^ vs. DDX3X^low^ cancer cells, respectively (Extended Data Fig. 6g). This *in silico* analysis of CCLE and DRIVE datasets demonstrates that DDX3X-low cancer cells share similar innate immune-activating signatures with ADAR1-dependent cancer cells, but also display the distinctive gene expression in T cell activation as well as immune cells activation.

### Inhibition of DDX3X suppresses breast tumor growth and enhances anti-tumor immunity

The expression of the type I IFN and chemokine (ex. CCL5) from the DDX3X-depleted cancer cells could activate effector T cells as well as other immune cells that could further prime T cells in the tumor. Also, the observation that DDX3X inhibition enhances the efficiency of the cancer cell’s antigen presentation through MHC class I prompted us to examine the tumor microenvironment. To investigate the impact of DDX3X inhibition on tumor immune microenvironment, we used the syngeneic mouse breast cancer model. We generated transplantable murine breast cancer cell line 4T1 that express doxycycline (Dox)-inducible DDX3X shRNAs, in which DDX3X level was considerably decreased after Dox treatment (Extended Data Fig. 7a). 1X10^5^ cells of 4T1 were inoculated into a mammary fat pad of immune-competent BALB/cJ female mouse. When tumors became palpable (20 to 25 mm^3^), randomized groups were treated with doxycycline foods to induce DDX3X knockdown. Tumor growth was monitored with calipers every three days. We found that inhibiting DDX3X substantially suppressed the 4T1 murine breast primary tumor growth, tumor volume, tumor weight, and metastatic spread to the lung in the BALB/cJ syngeneic mouse model (Fig. 7a-c and Extended Data Fig. 7b). The similar tumor growth inhibition by DDX3X depletion was also confirmed in the 4T1 tumor-bearing stable DDX3X KD in BALB/cJ female syngeneic mouse model (Extended Data Fig. 7c, d).

**Fig. 7:**
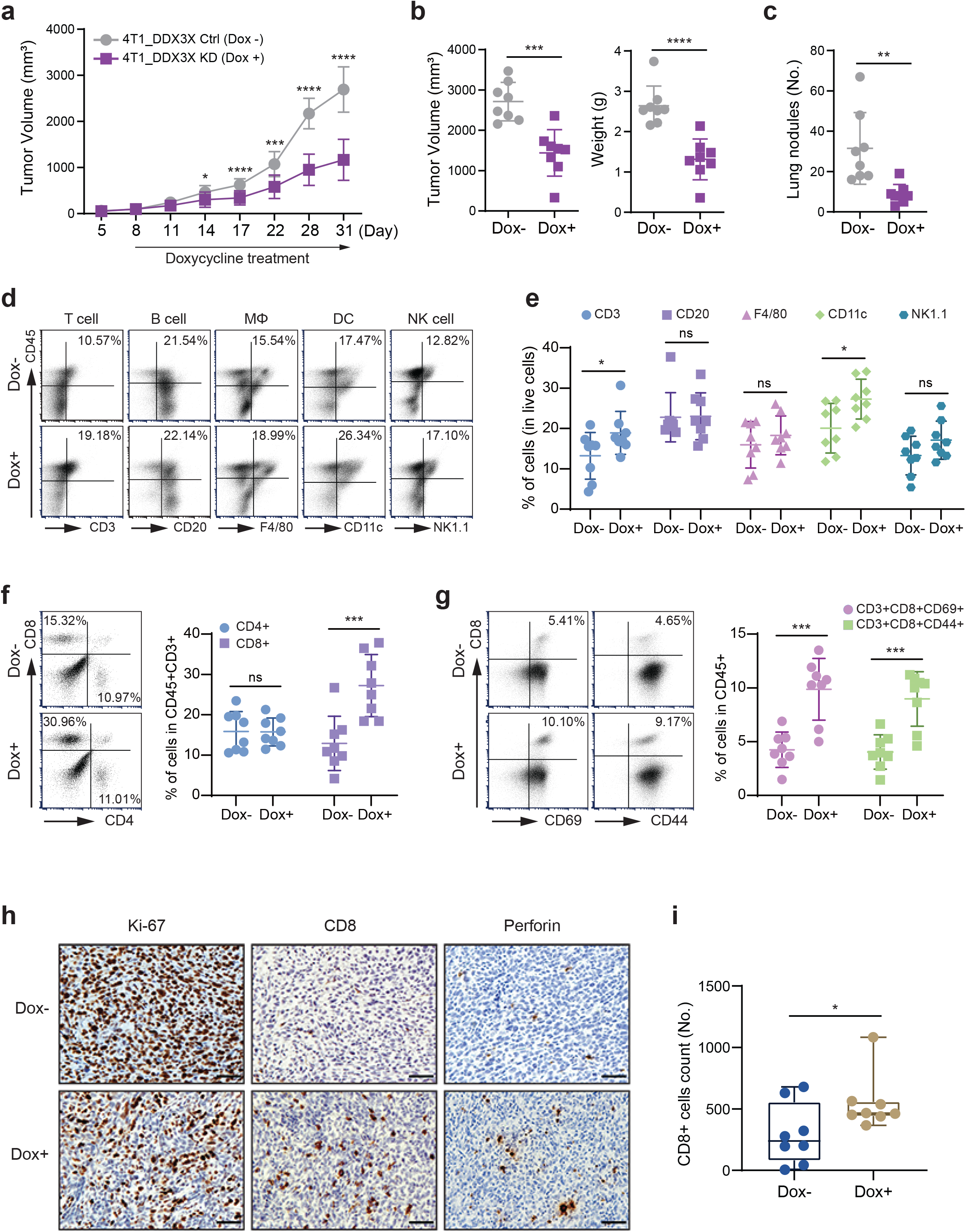
Inhibition of DDX3X suppresses breast tumor growth and enhances anti-tumor immunity. **a**, Tumor growth was monitored with calipers every three days. Mice bearing 4T1 tumors expressing doxycycline-inducible DDX3X shRNA were treated with (Dox+) or without doxycycline (Dox−) water. **b**, Tumor volumes and weights measured after dissection. **c**, Metastatic tumor nodules in lung counted after dissection. **d**, Representative dot plots showing each cell markers expression of CD45+ cells isolated from tumor. CD3+; T cell, CD20+; B cell, F4/80+; Macrophage, CD11c; dendritic cell, NK1.1; NK cell. **e**, Quantitative presentation of upper right quadrant from Fig. 6d. **f**, Representative dot plots showing CD4+ T cells or CD8+ T cells of CD45+ cells from tumors (left). Quantitative presentation of left dot plots (CD4+ T cells in lower right quadrant, CD8+ T cells in upper left quadrant). **g**, Representative dot plots showing activation profiles of CD8+ T cells of CD45+ cells isolated from tumors (left). Quantitative presentation of the upper right quadrants showing CD3+CD8+CD69+ or CD3+CD8+CD44+ T cells. **h**, Representative images of immunohistochemistry staining for CD8, perforin, and Ki67 in sections of tumors and quantitative data. Scare bars: 50 *μm*. **i**, Quantification of CD8 + cells in immunohistochemistry. Data are represented as mean ± SD. n=8 mice per group. Unpaired t-tests. \**P* < 0.05; \*\**P* < 0.01; \*\*\**P* < 0.001; \*\*\*\**P* < 0.0001; ns, not significant.

Flow cytometry analysis of the DDX3X-depleting 4T1 tumors revealed the significantly increased tumor infiltration of T cells and dendritic cells (DCs) (Fig. 7d, e). To further identify the T cell subsets, we assessed CD4+ helper and CD8+ cytotoxic T cells. The proportions of cytotoxic CD8+ T cells in the DDX3X-depleted tumor was increased, whereas the proportions of helper CD4+ T cell was present at a similar level between DDX3X-KD and -control tumor (Fig. 7f). Intratumoral CD8+ T cells in the DDX3X inducible-KD 4T1 tumors displayed the markedly increased expression of the effective cytotoxic CD8+ T cells markers, CD69 and CD44 (Fig. 7g). Similarly, significantly increased tumor infiltration of CD3+CD8+CD69+ T cells, CD3+CD8+CD44+ T cells, macrophage, and DCs was found in the DDX3X stable-KD 4T1 tumors (Extended Data Fig. 7e-g).

To obtain further insight within the tumors, we performed immunohistochemistry on the paraffin-embedded tumor sections. Consistent with the flow cytometry results, the increased CD8+ T cell infiltration was observed in the DDX3X-depleted tumors (Fig. 7h, i). Cytotoxic lymphocyte-secreted a pore-forming perforin, which shows tumor lytic functions, was also augmented in the DDX3X-depleted tumors (Fig. 7h). DDX3X-depleted tumors exhibited the decreased Ki67 proliferation marker (Fig. 7h). This *in vivo* result revealed that inhibition of DDX3X not only activates the tumor cell-intrinsic type I IFN response but also changes the tumor immune microenvironment resulting in increased tumor-infiltrating cytotoxic CD8+ T cells and DCs. Collectively, these data suggest that inhibiting DDX3X in tumors enhances the antigen presentation and the secretion of cytokines from cancer cells per se, which could induce enhanced anti-tumor immunity in the tumor microenvironment (Extended Data Fig. 7h).

## DISCUSSION

The findings in this study provide insights for the potential molecular mechanism of how DDX3X could regulate the cellular dsRNA homeostasis. Most endogenous dsRNAs are known to be located in the nucleus ^7^ or mitochondria ^15^, so that cells prevent these dsRNAs from being exposed to the cytosolic dsRNA sensors and triggering a potentially detrimental innate immune responses. The nuclear-cytoplasmic dsRNA distribution data indicates that a small fraction of dsRNAs is present in the cytoplasm in the homeostatic condition without inducing an immune response, but the depletion of DDX3X substantially increases the level of cytoplasmic dsRNAs (Fig. 5a). This suggests that unedited or partially edited genomic dsRNAs or mitochondrial dsRNAs could be transported to the cytoplasm ^24,62^ and DDX3X may be important to maintain the endogenous dsRNAs in the cytoplasm at a minimum level. There are several possibilities of how DDX3X could be involved in cellular dsRNA regulation. DDX3X could directly regulate the level or the stability of dsRNAs by unwinding the dsRNA structure. Alternatively, with the feature of a low complexity-intrinsically disordered region (IDR) in the N- and C-terminal sequences of DDX3X ^63–66^, it is possible that DDX3X sequesters dsRNAs within liquid-liquid phase-separations in the cytoplasm (e.g. small droplets or stress granules) until further degradation or modification of dsRNAs. Future questions that will need to be addressed are how ATP-, RNA-binding domains, and IDRs of DDX3X are associated with its dsRNA regulating function.

Our data also shows that dual depletion of DDX3X and ADAR1 produces a greater effect than single knockdown of each gene in activation of cytosolic dsRNA signaling pathway and innate immune response in the breast cancer cells (Fig. 5e-g). This suggests that DDX3X and ADAR1 may be acting independently on dsRNAs. Interestingly, depletion of DDX3X also inhibited ADAR1-mediated dsRNA editing activity when ADAR1 was depleted (Fig. 5h). Therefore, it is also possible that DDX3X could participate in ADAR1-mediated dsRNA editing. Because of the A-form structure of dsRNA, which includes a deep major groove ^67^, it is difficult for dsRNA-binding proteins like ADARs to recognize the dsRNA efficiently ^12^. Indeed, ADAR cannot bind and edit the dsRNA helices that are buried in a tertiary structure ^7^. As an RNA helicase, DDX3X could promote efficient ADAR-mediated dsRNA editing by unwinding the complex three-dimensional RNA structures and exposing dsRNA extension to ADAR1. Or DDX3X could facilitate other endo- or exonucleases for their processing or degradation of dsRNAs. To better understand the molecular mechanism of DDX3X-mediated dsRNA regulation, it will be important to identify the dsRNA species that are directly regulated by DDX3X in future study. Also, genome-wide sequencing analysis will be required to determine the direct impact of DDX3X on A-to-I dsRNA editing efficiency in the cells.

The present study additionally provides insight for a deeper understanding of the molecular mechanisms underlying the complex role of DDX3X in human innate immunity and cancer. DDX3X has been linked to viral replication and considered to promote antiviral response ^68–71^. In contrast, some viruses like HIV or HCV utilize DDX3X for their replication, suggesting that DDX3X could be a pro- or anti-viral factor ^43,72–74^. In particular, previous findings suggest DDX3X as a positive regulator of IFN production through the direct association with antiviral signaling pathways ^69,70,73,75^. In contrast to these findings, our study demonstrates that DDX3X prevents type I IFN production against endogenous dsRNAs by maintaining the homeostasis of endogenous dsRNAs. We note that the previous studies have been carried out in the cells after infection with virus, pathogens, or treatment with the viral mimicking synthetic dsRNAs, whereas the present findings are based on normal or malignant cells in the steady-state condition. Recently, Szappanos et.al showed that DDX3X is an essential for innate immunity against the infection of pathogenic bacteria, but DDX3X is also important for hematopoiesis and keeping maintenance of immune cells in mice, with the most profound effects observed in lymphocytes and splenic NK cells ^76^. Most recently, Samir et.al showed a role for DDX3X in driving NLRP3 inflammasome and stress granule assembly. They showed that DDX3X interacts with NLRP3 and is required for NLRP3 inflammasome activation in bone-marrow-derived macrophages stimulated with lipopolysaccharide ^77^. Given the previous and our current findings, we propose that DDX3X may have a “dual effect” on immune activity in humans (anti-autoimmunity and anti-viral activity). Although more detailed studies are required to mechanistically define antiviral activity and anti-autoimmunity against cellular dsRNAs of DDX3X, the current study proposes the complex role of DDX3X in the dsRNA biology network and antiviral immunity. In the future, it will be important to investigate how different RNA viruses utilize this potential dual role of DDX3X for their replication. Our conditional DDX3X knockout mouse could be a useful *in vivo* tool to systematically define the role of DDX3X.

Loss of ADAR1 increases cellular dsRNAs that trigger the type I IFN responses in the various tumor cells and affording increased sensitivity to immunotherapy ^17,78,79^. Recently, Liu et al. reported that cancer cells have chronic type I IFN activation and increased ISG signature triggered by a STING-dependent pathway, rendering them sensitive to ADAR1 loss ^58^. The malignant cells may have elevated levels of dsRNAs due to the loss of suppressive epigenetic modifications in repetitive elements, genomic instability, or mitochondrial damage due to metabolic stress, which would increase dsRNA burden in the cells ^12,80^. Therefore, cancer cells with the high dsRNAs load could rely more on the active cellular dsRNA regulatory mechanism to avoid the activation of innate immune signaling as well as to escape them from immune surveillance. CCLE analysis results suggest that DDX3X-dsRNAs-type I IFN response axis is not restricted to the specific type of cancer such as breast cancer, but rather depends on the endogenous dsRNAs expression burden or intact innate immune systems of the cancer cells. In GO analysis of DEGs, both ADAR and DDX3X are related to the strong upregulation of antiviral immune signatures including type I interferon signaling pathway and innate immune response, suggesting that DDX3X and ADAR1 are possibly functionally linked to each other in various cancer cells. Notably, the DDX3X expression level is inversely correlated with the immune cell activation, including T cell activation, lymphocyte proliferation, and regulation of the immune effector process (Fig. 6e). This suggests that the diverse dsRNA regulatory mechanism could induce a different impact on the tumor immune microenvironment.

Despite an established benefit of type I IFN in cancer therapy, the exogenous administration of type I IFN has shown systemic adverse effects and at best modest antitumor efficacy, which has reduced its use ^81^. Recent findings of endogenous nucleic acid-induced tumor cell-intrinsic type I IFN pathway gives new insights for enhancing the therapeutic effect of type I IFN ^81,82^. Tumor cell-intrinsic type I IFN response has been achieved through the intracellular dsRNA transcription by using a CDK4/6 inhibitor ^16^ or DNA methylation inhibitors ^18,26,83^. Also, cellular dsRNA accumulation has been induced at the post-transcriptional level by inhibiting ADAR1 ^17^, LSD1 ^14^, or PNPT1 ^15^. Our data shows that inhibiting DDX3X enhances cancer cell-intrinsic type I IFN response, antigen presentation on the cancer cells, and the antitumor activity in tumors. In particular, inhibiting DDX3X increased not only tumor-infiltrated cytotoxic CD8+T cells, but also DCs. CD8+ T cells recognize tumor-associated antigens presented on MHC I by their expressed T cell receptor (TCR) to remove tumor cells and, DCs are required to cross-present exogenous tumor-associated antigens onto MHC I to prime CD8+ T cells ^84^. Furthermore, DDX3X depleted-OVA expressing cancer cells showed the increased ability of antigen presentation onto MHC I. These data show that inhibiting DDX3X could enhance the elimination of cancer cells by CD8+ T cells directly and DCs indirectly. Increased tumor cell antigen presentation coupled with anti-tumor T-cell responses suggests that immune checkpoint blockade might further enhance the efficacy of DDX3X inhibition.

In conclusion, the present study identifies a role of DDX3X in regulating the endogenous dsRNAs homeostasis in human and mouse malignant cells as well as MEFs. DDX3X maintains the cytosolic endogenous dsRNAs at a low level, but the loss of DDX3X leads to the aberrant accumulation of cellular dsRNAs, which activates cytoplasmic dsRNA sensors and triggers type I interferon signaling in the breast cancer cells. We further show that inhibiting DDX3X restores cancer immunity and enhances the antitumor activity by inducing the dsRNA-derived type I IFN response in breast tumors, which may lead to novel agents targeting DDX3X for combinatory immunotherapy.

## ACKNOWLEDGEMENTS

We thank Michael B. Atkins and Xiongbin Lu for a critical reading of the manuscript. We thank Marta Catalfamo, Astrid Haase, and Pavol Genzor for a helpful scientific discussion. We thank Marie Öhman for the kind gift of ADAR1 editing vectors. We thank Charles G. Drake and Mark Smyth for sharing B16-OVA cells. We thank Jing Wang and Xiaogang Zhong for helping the RNA sequencing analysis. This work was in part supported by an NIH/NCI Pathway to Independence Award (K99/R00 CA197487).

## AUTHOR CONTRIBUTIONS

Conceptualization, C.H and J.C.; Methodology, M.S.C., and Y.S.; Investigation, H.C., J.K., J.S., and C.H.; Writing – Original Draft, H.C. and J.K.; Writing – Review & Editing, C.H. J.C.; Funding Acquisition, C.H.; Supervision, C.H.

## EXPERIMENTAL PROCEDURES

### Cell culture and generation of stable cell lines

MCF7, MDA-MB-453, MDA-MB-231, A375, B16F10, 4T1, E0771, and HEK293T cell lines were obtained from the American Type Culture Collection and cultured under standard conditions specified by the manufacturer. B16-OVA cells were kindly provided by Dr. Charles G. Drake (Columbia University Medical Center, NY, USA). The cells were maintained in DMEM medium supplemented with 10% fetal bovine serum, 1% penicillin/streptomycin, and 250 μg/ml G418 (Invitrogen) at 37°C with 5% CO_2_. All the cell lines were tested negative for mycoplasma using the Mycoplasma Detection kit (Lonza). For generation of stable DDX3X knockdown cell lines, lentiviral pGIPZ, inducible TRIPZ, and SMARTvector (Inducible Mouse Ddx3x shRNA) were obtained from Dharmacon. The following sequences are targeted: GIPZ Lentiviral Human DDX3X shRNA #1_Clone Id_V2LHS_228965: 5’-TAAATCTGACTCAAGATGG-3’, GIPZ Lentiviral Human DDX3X shRNA #2_Clone Id_V3LHS_301003: 5’-GTACTGCCAACTCTCTCGT-3’, GIPZ negative (non-targeting or non-silencing) shRNA control_Catalog ID_RHS4346, TRIPZ Inducible Lentiviral shRNA Human DDX3X #1_Clone Id_V3THS_301003: 5’-GTACTGCCAACTCTCTCGT-3’, TRIPZ Inducible Lentiviral shRNA Human DDX3X #2_Clone Id_V2THS_228965: 5’-TAAATCTGACTCAAGATGG-3’, SMART vector Inducible Mouse Ddx3x PGK-TurboRFP shRNA #1_Clone Id_ V3SM11253-232377132: 5’-TCCCTCTTGAATCACCCCG-3’, SMARTvector Inducible Mouse Ddx3x PGK-TurboRFP shRNA #2_Clone Id_V3SM11253-232987995: 5’-TGCACTGCCAATTCTCTCG-3’. Recombinant lentiviral particles were produced using a protocol provided by the manufacturer (Addgene). In brief, 2 μg of shRNA-encording vector DNA, 1.5 μg of psPAX2 (packaging vector) and pMD2.G (VSVG envelope vector) vectors were transfected into HEK293T cells in 94 mm^2^ dish using TransIT-LT1 Transfection Reagent (Mirus). The supernatants containing virus particles was collected at 72 hours after transfection. Filtered viral supernatants added in the growth medium in the presence of polybrene (8 mg/ml). To establish stale KD cell lines, cells were selected 48 hours after viral infection using 2 mg/ml of puromycin and collected single colonies and propagated. Knockdown efficiency was validated by western blotting (protein) and qRT-PCR (mRNA). The IFIH1 gene was deleted using a MDA5 (IFIH1) Human Gene Knockout Kit (OriGene, KN415661) according to the manufacturer’s instructions with target sequences (5’-CTGGATGTACATTTTCACCC-3’).

### Immunohistochemistry and immunofluorescence

For immunohistochemistry, paraffin embedded tumor samples and sections were prepared as previously described ^85^. Samples were incubated with primary antibodies which were CD8 (#98941, Cell Signaling Technology), Ki-67 (#12202, ell Signaling Technology), and Perforin (#31647, Cell Signaling Technology) for overnight at 4°C. Sections were incubated with biotinylated goat anti-rabbit IgG (PK-4000, Vector laboratories) followed by washing three times with TBS-T (0.05% Tween-20). After washing, sections were incubated with VECTASTAIN ABC Reagent (PK-4000, Vector laboratories) for 30 minutes and then developed using a DAB peroxidase substrate kit (ImmPACT DAB, SK-4105, Vector laboratories). Sections were counterstained with hematoxylin (3801570, Leica). For immunofluorescence, cells were plated on glass coverslips. Cells were washed with PBS for two times and fixed with 4% (v/v) formaldehyde in PBS at room temperature for 15 minutes and permeabilized with 0.25% Triton-X100 in PBS for 20 minutes at room temperature. J2 antibody was diluted in PBS (1: 100) and incubated at 4°C for overnight. Cells were washed with PBS for three times and incubated with rat anti-mouse IgG2a conjugated with APC (407109, BioLegend) for 1 hour at room temperature. Imaging was performed with a Leica SP8 confocal microscope. For nuclease treatment, cells were treated with enzymes at 37°C for 30 minutes before fix the cells. Following enzymes were used: RNase I (50 U/ml), RNase A (10 μg/ml), RNase III (20 U/ml) in PBS containing 5 mM MgCl_2_.

### Western blot

Cells were washed with ice-cold PBS twice and aspirated residual medium completely. Cell lysis buffer (#9803, Cell Signaling Technology, composition: 20 mM Tris-HCl (pH 7.5), 150 mM NaCl, 1 mM Na_2_EDTA, 1 mM EGTA, 1% Triton, 2.5 mM sodium pyrophosphate, 1 mM β-glycerophosphate, 1 mM Na3VO4, 1 μg/ml leupeptin, 1mM PMSF, and protease inhibitor cocktail (87785, Thermo Fisher)) was added to cell dishes and scraped on ice. Cell lysates were keep on ice for 30 minutes and centrifuged at 13,200 rpm for 10 minutes at 4°C. Protein concentrations were measured by Bio-Rad protein assay (#5000006, Bio-Rad) and equal volume and quantity of protein samples were made by addition of 4x Laemmli Sample Buffer (#1610747, Bio-Rad) containing 2-merchatoethanol (#1610710, Bio-Rad) and boiling at 95°C for 10 minutes. Protein samples were subjected to SDS-PAGE and transferred to polyvinylidene difluoride membranes (#1620177, Bio-Rad). The membranes were blocked with 5% skim milk in TBS-T (0.1% Tween-20) for 30 minutes at room temperature and incubated with indicated primary antibodies at 4°C overnight. On the next day, membranes were washed and incubated with appropriate peroxidase-conjugated secondary antibodies for 1 hour at room temperature. Following several washes, ECL (WBKLS0500, Millipore or #1705061, Bio-Rad) was applied for membrane development. Following antibodies were used: rabbit anti-DDX3 (A300-474A, Bethyl Laboratories), mouse anti-DDX3 (sc-365768, Santa Cruz), rabbit anti-DDX1 (A300-521A, Bethyl Laboratories), rabbit anti-DDX5 (A300-523A, Bethyl Laboratories), rabbit anti-ADAR1 (A300-884A, Bethyl Laboratories), mouse anti-ADAR1 (sc-271854, Santa Cruz), mouse anti-β-Actin (sc-4777, Santa Cruz), rabbit anti-phospho-Stat1 (Tyr701) (#9167, Cell Signaling Technology), rabbit anti-Stat1 (#14994, Cell Signaling Technology), rabbit anti-phospho-Stat2 (Tyr690) (#4441, Cell Signaling Technology), rabbit anti-Stat2 (#72604, Cell Signaling Technology), rabbit anti-RIG-I (#4200, Cell Signaling Technology), rabbit anti-MDA-5 (#5321, Cell Signaling Technology), rabbit anti-OAS1 (#14498, Cell Signaling Technology), rabbit anti-phospho-PKR (Thr451) (#07-886, Millipore), mouse anti PKR (sc-6282, Santa Cruz), rabbit anti-phospho-eIF2a (Ser51) (#9721, Cell Signaling Technology), rabbit anti-eIF2α (#9722, Cell Signaling Technology), rabbit anti-cGAS (#15102, Cell Signaling Technology), rabbit anti-phospho-STING (Ser366) (#85735, Cell Signaling Technology), rabbit anti-STING (#13647, Cell Signaling Technology), rabbit anti-phospho-NF-kB p65 (Ser536) (#3033, Cell Signaling Technology), rabbit anti-NF-kB p65 (#8242, Cell Signaling Technology), rabbit anti-phospho-IRF-3 (Ser396) (#29047, Cell Signaling Technology), rabbit anti-phospho-IRF-7 (Ser477) (#12390, Cell Signaling Technology), rabbit anti-Toll-like Receptor 3 (#6961, Cell Signaling Technology), mouse anti-Lamin A/C (sc-376248, Santa Cruz), mouse anti-alpha tubulin (ab7291, Abcam), rabbit anti-Drosha (A301-886A, Bethyl Laboratories), mouse anti-FLAG M2 (F1804, Sigma), mouse anti-IFNγRα (sc-12755, Santa Cruz), mouse anti-IFNαβRα (sc-7391, Santa Cruz).

### Immunoprecipitation

Cells were lysed in immunoprecipitation buffer (1% NP-40, 50 mM Tris-HCl, 500 mM NaCl and 5 mM EDTA) containing protease inhibitor cocktails (87785, Thermo Fisher) on ice for 30 minutes. Cell lysates (800 μg) were incubated with 3 μg of antibodies or normal IgG at 4°C overnight with rotary agitation. For immunoprecipitation of DDX3X or ADAR1, anti-DDX3X (sc-365768, Santa Cruz) or anti-ADAR1 (sc-271854, Santa Cruz) was used, respectively. Protein G agarose beads (11243233001, Roche) were added to the lysates and incubated for additional 4 hours at 4°C with rotary agitation. The IP Beads were washed in immunoprecipitation buffer three times for 10 minutes each and completely removed residual buffer and boiled in SDS loading buffer for 10 minutes at 95°C for western blot analysis. For J2 immunoprecipitation, cells were fixed with 1 % formaldehyde prior to lysis cells. Immunoprecipitation was performed with J2 antibody (10010200, Scicons) and normal IgG, respectively and coupled to the protein G agarose beads.

### J2 dot blot and northwestern blot

Total cellular RNA or the RNA of cytoplasmic and nucleus fractions were extracted using the manufacturer’s protocol of TRIzol reagent (15596026, Thermo Scientific) or TRIzol LS Reagent (10296010, Thermo Scientific), respectively. RNA concentration was measured by the Nonodrop. For dot blot, purified RNA was dotted on PVDF membrane following pre-dotted site using methanol. The membrane was dried on table for overnight. For northwestern blot, purified RNA was electrophoresed in non-denaturing 2% agarose TBE gel and then gel was blotted onto an Hybound-N nylon membrane (RPN303N, GE Healthcare) by semidry electroblotting in 0.5× TBE buffer for 2 hours at 200mA. The membrane was blocked in 5% skim milk in TBS-T (0.1% Tween-20). J2 antibody was incubated for overnight at 4°C and then secondary antibody was incubated for 1 hour at room temperature. ECL was applied for the membrane visualization (WBKLS0500, Millipore or #1705061, Bio-Rad).

### qRT-PCR

Total cellular RNA or the RNA of cytoplasmic and nucleus fractions were extracted using the manufacturer’s protocol of TRIzol reagent (15596026, Thermo Scientific) or TRIzol LS Reagent (10296010, Thermo Scientific), respectively. RNA concentration was measured by the Nonodrop and 1 μg of total RNA was used to generate cDNA with the High Capacity cDNA Reverse Transcription Kits (4368814, Thermo Scientific). Quantitative reverse transcription PCR (qRT-PCR) of *DDX3X*, *IFI44L*, *IFIT2*, *IFIT3*, *ISG15*, *STAT1*, *OAS1*, *OAS2*, *MX1*, *TAP1*, *TAP2*, *PSMB8*, *HLA-A*, *HLA-B*, *HLA-C*, *HLA-DRA*, *RIG-I*, *MDA5*, and *IFNB1* was performed using TaqMan Fast Advanced Master Mix (4444557, Applied Biosystems). qRT-PCR of *envK1-6*, *envV1*, *MER21C*, *MER57B1*, *ERVF*, *ERVFb1*, *ERV9-1*, *MATA10*, *MLT2B4*, *ENV-MER34*, and *ERV-Fb1* was performed using SYBR Green master mix (A25742, Applied Biosystems). All qRT-PCR was performed on a QuantStudio 3 Real-Time PCR system and software (Applied Biosystems). *GAPDH* or *18s* were used as reference genes. All qRT-PCR assays were carried out in triplicate and then repeated with new cDNA synthesis. Primer information is summarized in Supplemental Table 2.

### Animal experiments

Female BALB/cJ mice (6–8 weeks old) were purchased from Jackson Laboratories. All studies were approved and supervised by the Institutional Animal Care and Use Committee at Georgetown University. To generate syngeneic mouse mammary tumor model, 1×10^5^ 4T1-inducible DDX3X knockdown cells were implanted into mammary fat pad after mixed 1:1 by volume with matrigel (Corning). Before inducing of DDX3X knockdown, mice were divided into control and DDX3X knockdown groups of equal average tumor volume. Doxycycline water (5% sucrose with 2 mg/ml of doxycycline (Sigma) was provided to DDX3X knockdown group to induce knockdown of DDX3X from 1 week after cell injection. The doxycycline water was changed every other day. Tumor size was measured every 3-5 days by caliper. The tumor volume calculated using the formula: Volume = (Width^2^ × Length)/2.

### Flow cytometry

For tumor infiltrating leukocyte flow cytometry, tumors (0.5 g) were mechanically disrupted by chopping and then chemically digested using Tumor Dissociation Kit (130-096-730, Miltenyi Biotec) and gentleMACS Dissociator (130-096-427, Miltenyi Biotec) according to the manufacturer’s instructions. Isolated tumor cells were lysed red blood cells (10-548E, Lonza) and blocked Fc receptors with anti-CD16/32 (101319, BioLegend) for 20 minutes on ice and then stained with appropriate antibodies for 1 hour on ice. Before flow cytometry analysis, cells were stained with Zombie NIR Fixable Viability Kit (423105, BioLegend) to distinguish live or dead cells. For OVA or HLA class expression on cell surface, cells were detached with 2 mM EDTA in PBS and then washed twice using PBS before staining. Cells were stained with appropriate antibodies for 30 minutes on ice and then washed twice before flow cytometry analysis. The following antibodies were used: APC conjugated anti-mouse H-2Kb bound to SIINFEKL (141605, BioLegend) and mouse IgG1, κ Isotype control (555751, BD Pharmingen), APC conjugated anti-human HLA-DR (307609, BioLegend) and mouse IgG2a isotype control (407109, BioLegend), and APC conjugated Anti-Human HLA-ABC (562006, BD Pharmingen) and mouse IgG1, κ Isotype control (555751, BD Pharmingen). For J2 flow cytometry, cells were detached with 2 mM EDTA in PBS and then washed twice using PBS. Cells were fixed with 1x fixation buffer (424401, BioLegend) for 20 minutes at room temperature. After washing, cells were permeabilized with 0.1% Triton-X-100 in PBS for 15 minutes followed by incubation in 1% BSA in PBS for 30 minutes. Cells were stained with J2 antibody (10010200, Scicons) or mouse IgG2a isotype control (401501, BioLegend) for 1 hour at room temperature followed by anti-mouse IgG2 conjugated with APC was stained for secondary antibody. Flow cytometry were performed with a FACSCalibur (BD Biosciences) and data were analyzed using FCS express 6.

### CCLE data analysis

Gene expression and gene effect data were obtained from the CCLE and DeMap portal (http://doi:10.6084/m9.figshare.11384241.v2). Cell lines with fibroblast and hematopoietic lineage expression profile were excluded from the analysis. DDX3X high expressing (DDX3X^hi^) cells and DDX3X low expressing (DDX3X^low^) cells were selected by DDX3X expression level (top 25% and bottom 25%, respectively). ADAR1 dependency data was downloaded from DeMap portal. Gene effect score is obtained by large scale RNAi screening. A lower score means that a gene is more likely to be dependent in a given cell line. We used gene dependency score as a reverse value of gene effect score. ADAR1 dependent- (ADAR1^dep^) and independent- (ADAR1^idp^) groups were divided by ADAR1 gene dependency (top 25% and bottom 25%, respectively). RNA-seq data from selected groups was normalized with the voom method and differential expression determined by limma ^86^. MHC core scores were calculated with mean absolute deviation modified Z-score mRNA expression data of in CCLE. The score was defined as the mean Z-score of all MHC class genes in each group. GSEA analysis was performed using fgsea function ^87^ and KEGG and GO term enrichment analysis on expression data was conducted using clusterProfiler function from R Bioconductor package ^88^. To select immune-related genes, differentially expressed genes in each group were compared with innateDB (https://www.innatedb.com) ^89^. Differentially expressed immune-related gene lists were subjected as an input for GO enrichment analysis

