## Supplemental information for "Inhibiting DDX3X triggers tumor-intrinsic type I interferon response and enhances anti-tumor immunity"

### Extended Information

- Extended Figures 1-7
- Extended Figures legends
- Extended Experimental Procedures
- Extended References
- Supplementary Table 1-Related to Figure 1A (Gene list of differentially expressed genes)
- Supplementary Table 2-Primer information for qRT-PCR and TASA-TD PCR

***Extended. Fig.1***

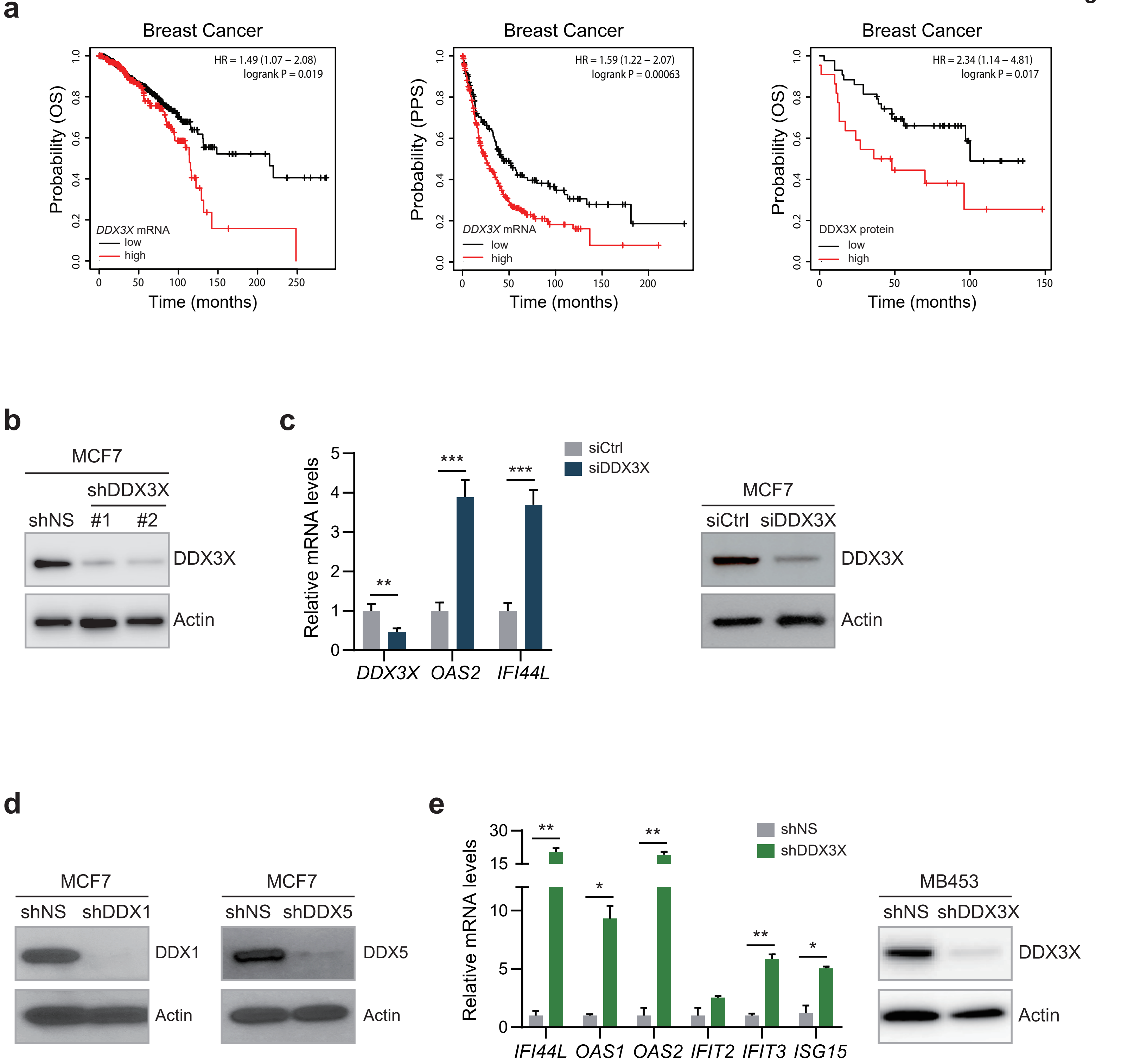

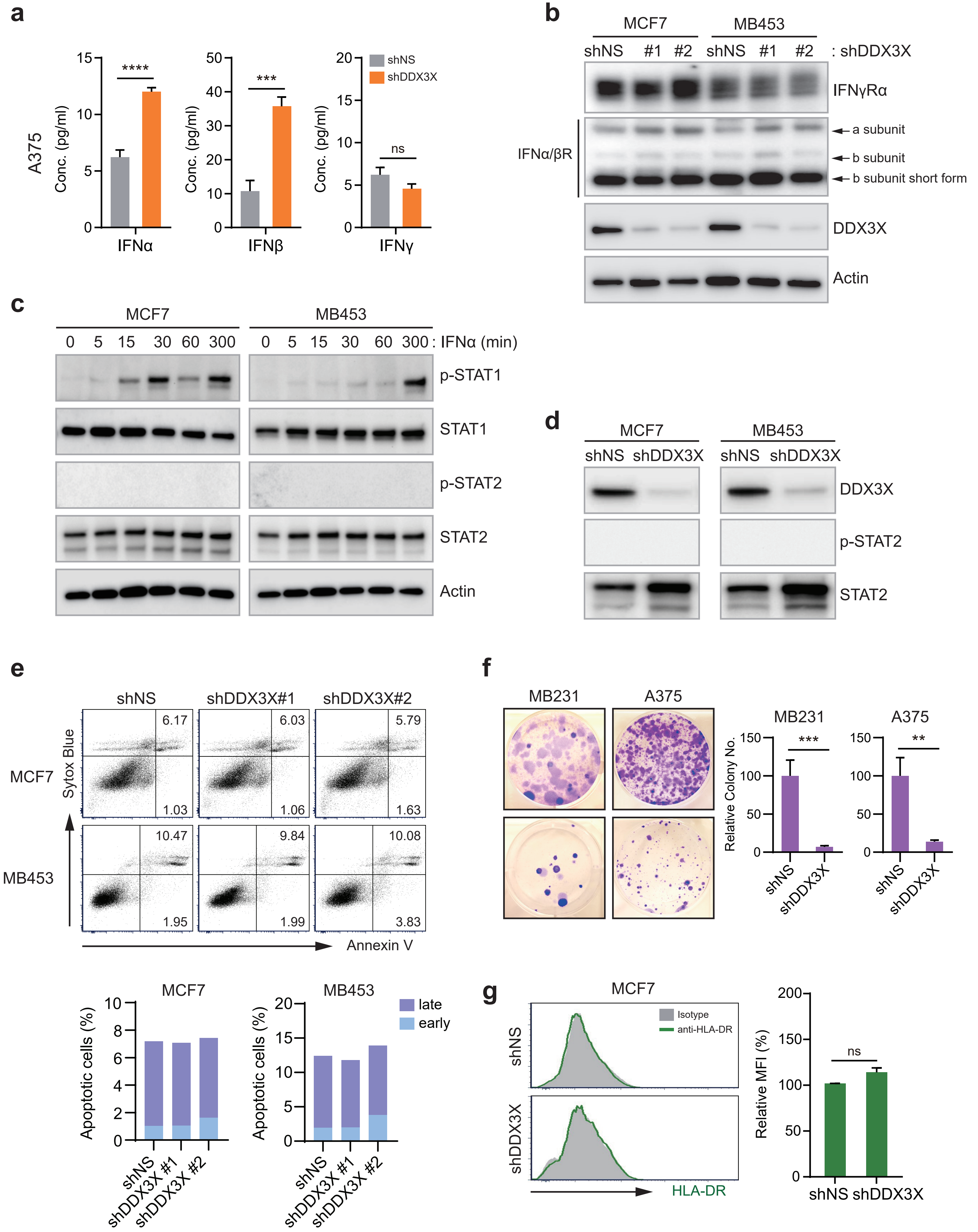

a
A375
Conc. (pg/ml)
****
***
ns
shNS
shDDX3X
IFNα
IFNβ
IFNγ
b
MCF7
MB453
shNS #1 #2 shNS #1 #2 : shDDX3X
IFNγRα
IFNα/βR
a subunit
b subunit
b subunit short form
DDX3X
Actin
c
MCF7
MB453
0 5 15 30 60 300 : IFNα (min)
p-STAT1
STAT1
p-STAT2
STAT2
Actin
d
MCF7
MB453
shNS shDDX3X
DDX3X
p-STAT2
STAT2
e
shNS
shDDX3X#1
shDDX3X#2
MCF7
MB453
Sytox Blue
Annexin V
6.17
6.03
5.79
1.03
1.06
1.63
10.47
9.84
10.08
1.95
1.99
3.83
Apoptotic cells (%)
late
early
shNS
shDDX3X #1
shDDX3X #2
f
MB231
A375
Relative Colony No.
***
**
shNS
shDDX3X
g
MCF7
Isotype
anti-HLA-DR
shNS
shDDX3X
HLA-DR
Relative MFI (%)
ns

***Extended Fig. 3***

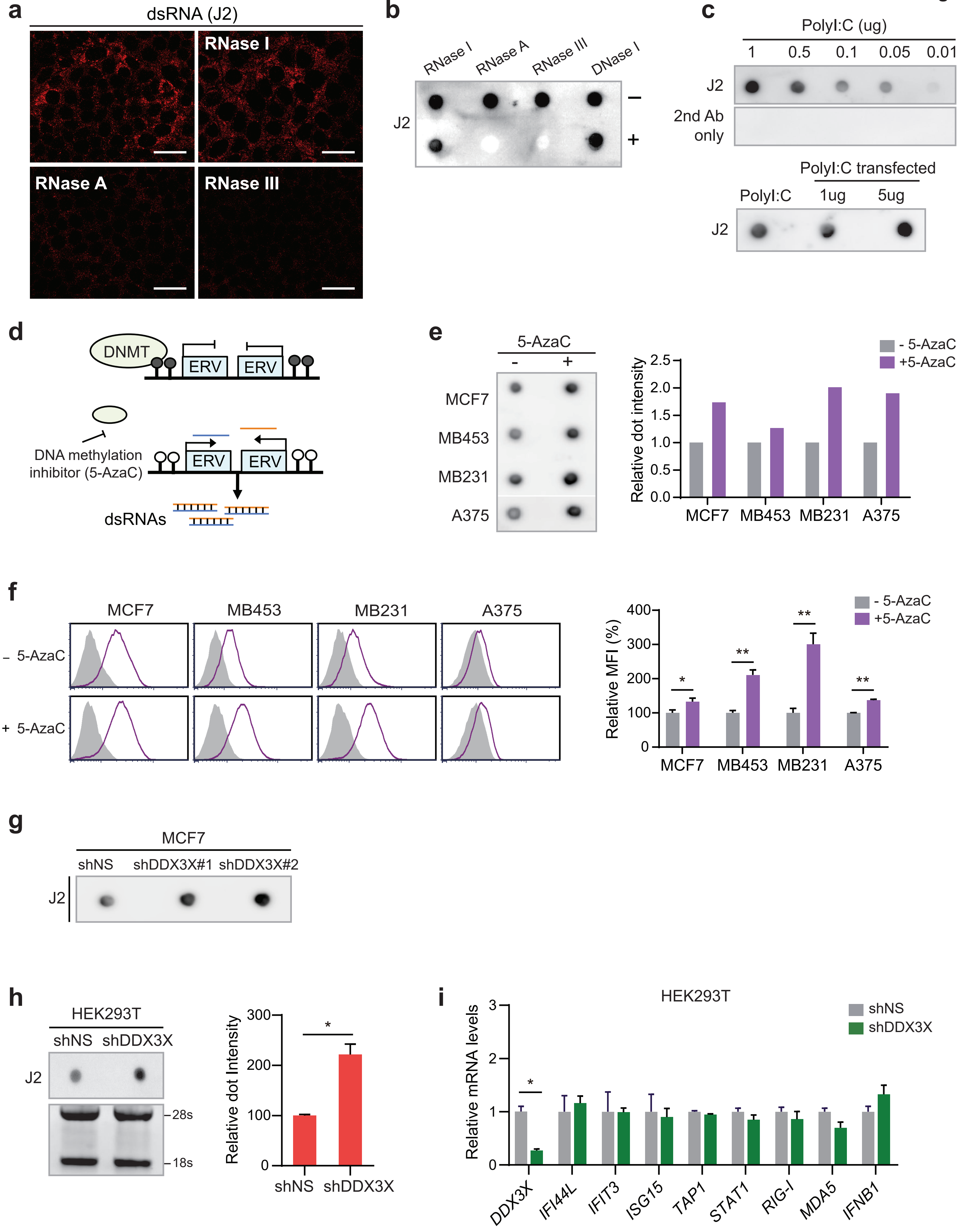

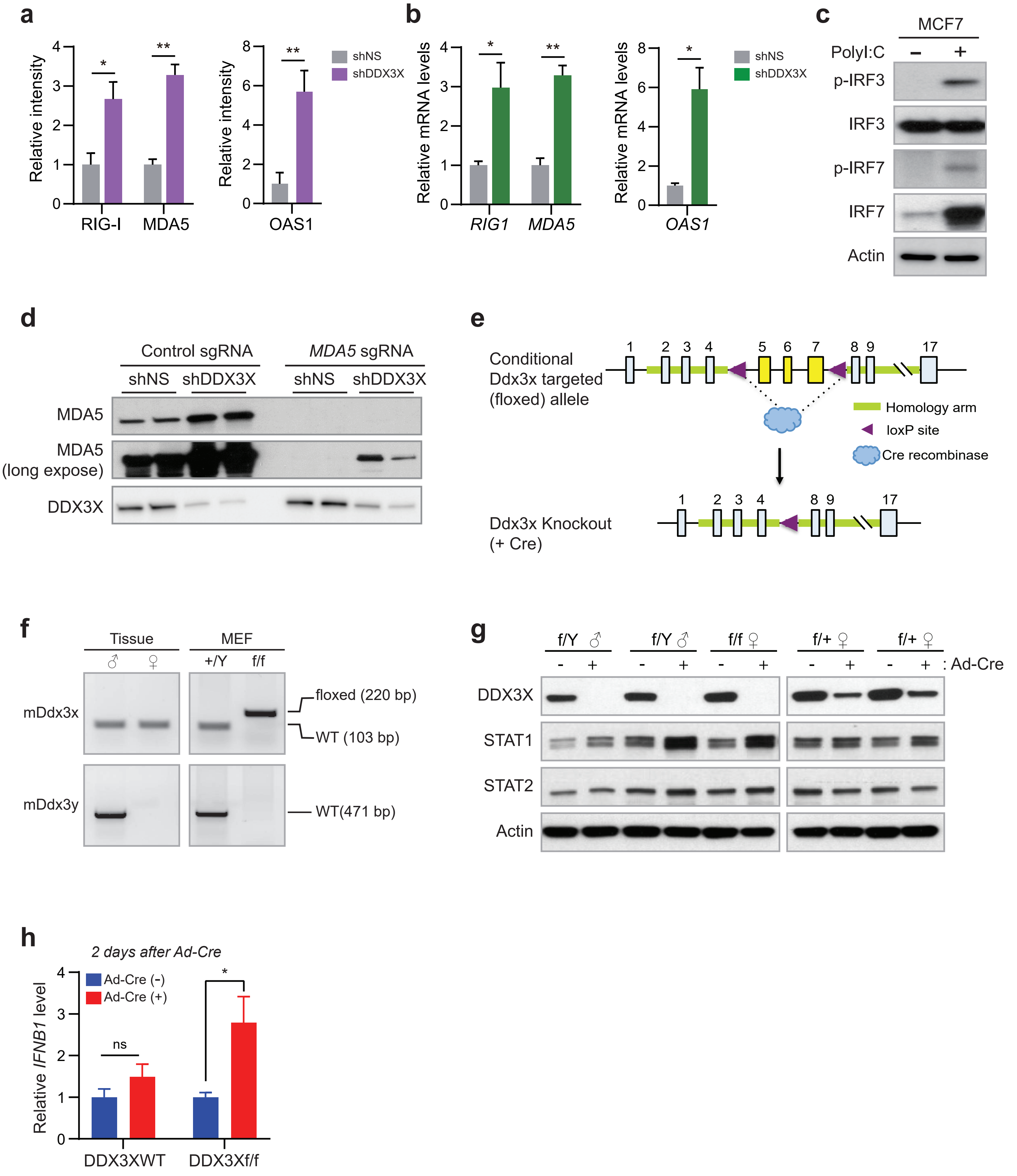
a
Relative intensity
RIG-I
MDA5
OAS1
shNS
shDDX3X
b
Relative mRNA levels
RIG1
MDA5
OAS1
shNS
shDDX3X
c
MCF7
PolyI:C
p-IRF3
IRF3
p-IRF7
IRF7
Actin
d
Control sgRNA
MDA5 sgRNA
shNS
shDDX3X
MDA5
MDA5 (long expose)
DDX3X
e
Conditional Ddx3x targeted (floxed) allele
Homology arm
loxP site
Cre recombinase
Ddx3x Knockout (+ Cre)
f
Tissue
MEF
+/Y
f/f
mDdx3x
mDdx3y
floxed (220 bp)
WT (103 bp)
WT(471 bp)
g
f/Y ♂
f/f ♀
f/+ ♀
: Ad-Cre
DDX3X
STAT1
STAT2
Actin
h
2 days after Ad-Cre
Relative IFNB1 level
Ad-Cre (-)
Ad-Cre (+)
ns
DDX3XWT
DDX3Xf/f

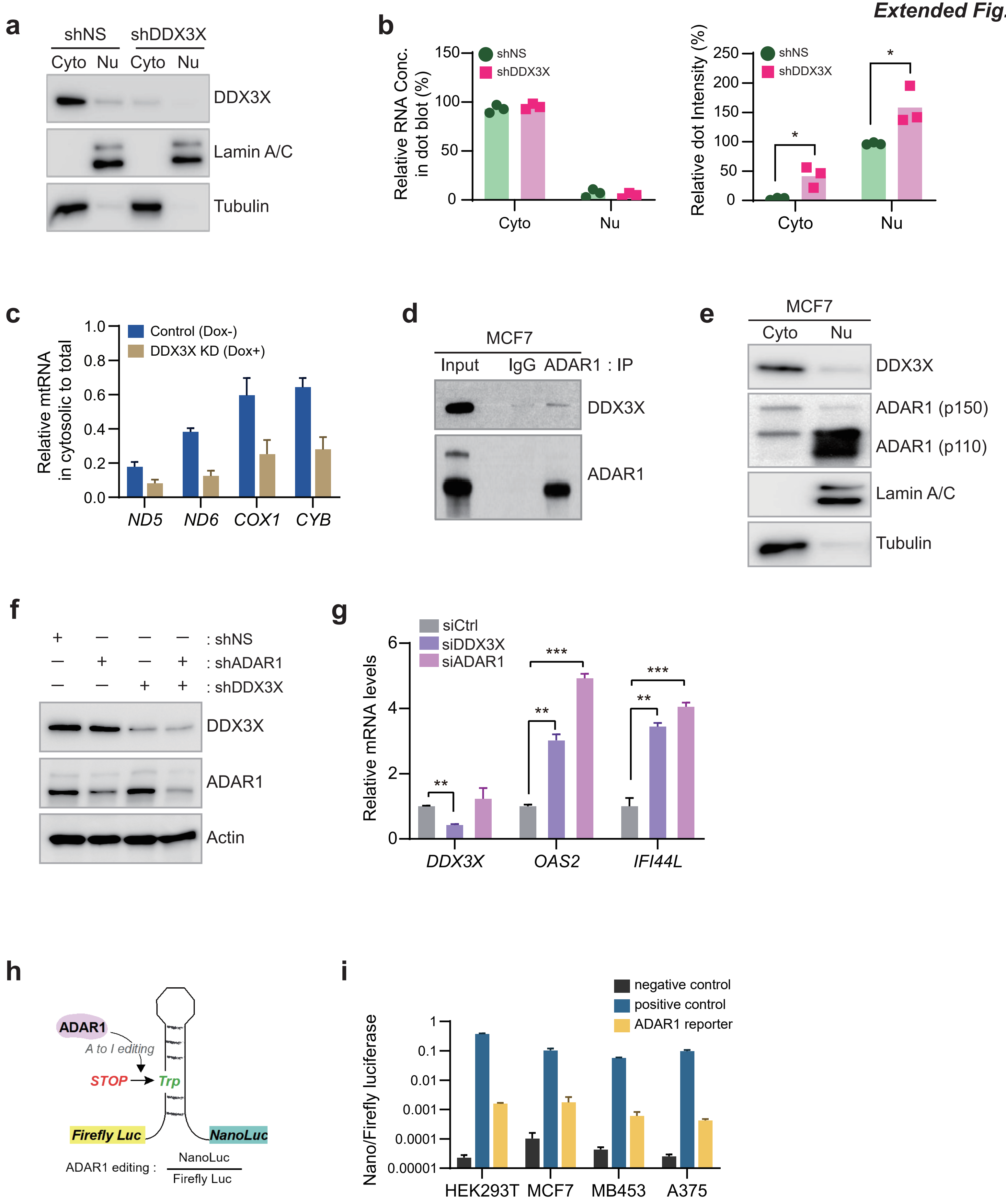
a
shNS
shDDX3X
Cyto Nu Cyto Nu
DDX3X
Lamin A/C
Tubulin
b
Relative RNA Conc. in dot blot (%)
shNS
shDDX3X
Cyto
Nu
Relative dot Intensity (%)
*
c
Relative mtRNA in cytosolic to total
Control (Dox-)
DDX3X KD (Dox+)
ND5
ND6
COX1
CYB
d
MCF7
Input IgG ADAR1 : IP
DDX3X
ADAR1
e
MCF7
Cyto Nu
DDX3X
ADAR1 (p150)
ADAR1 (p110)
Lamin A/C
Tubulin
f
: shNS
: shADAR1
: shDDX3X
DDX3X
ADAR1
Actin
g
siCtrl
siDDX3X
siADAR1
Relative mRNA levels
**
***
DDX3X
OAS2
IFI44L
h
ADAR1
A to I editing
STOP
Trp
Firefly Luc
NanoLuc
ADAR1 editing :
NanoLuc
Firefly Luc
i
negative control
positive control
ADAR1 reporter
Nano/Firefly luciferase
HEK293T
MCF7
MB453
A375

### Extended Fig. 6

**a**

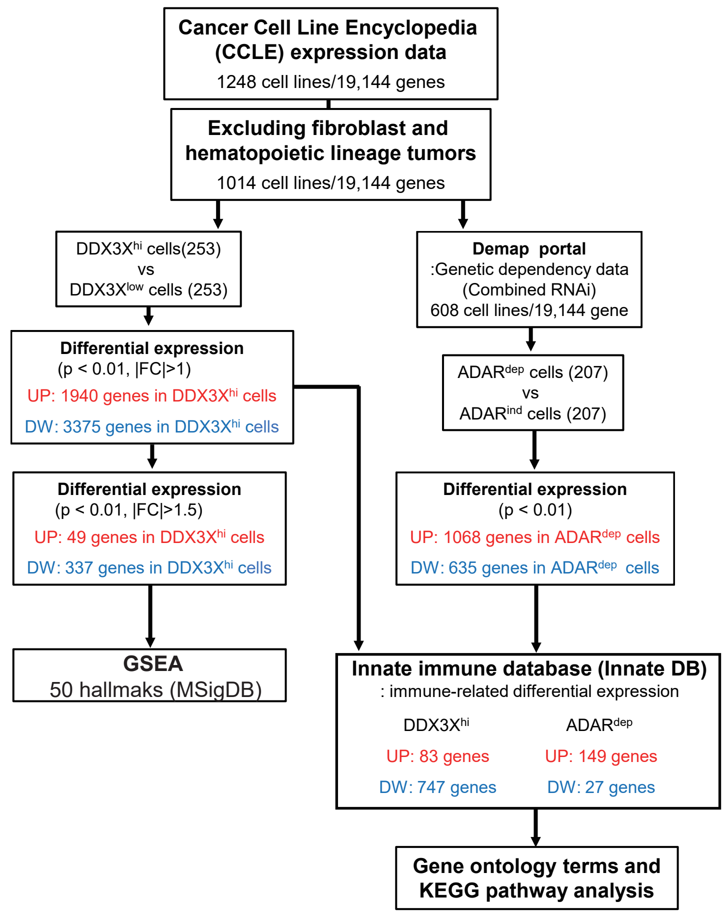


**b**

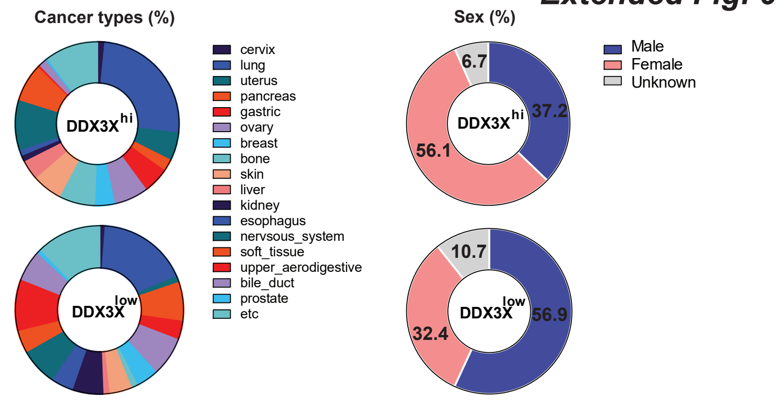


**d**

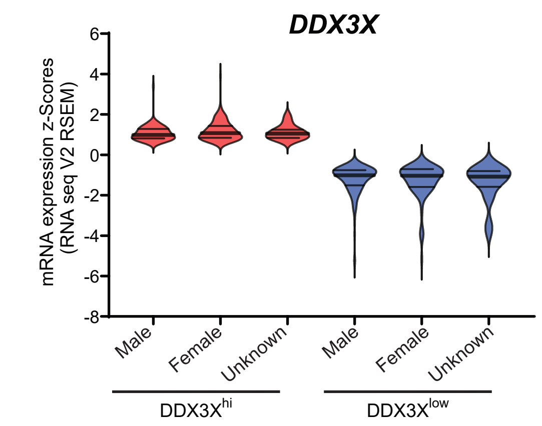


**e**

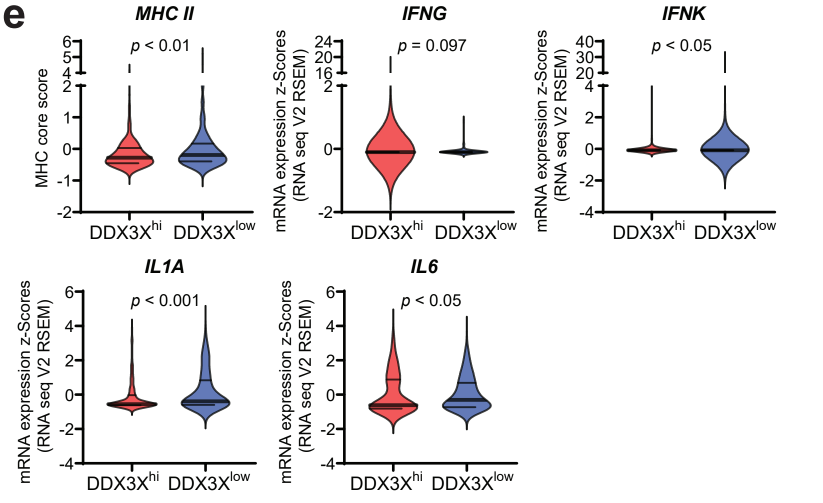


**d**

| 6-gene MHC class I | | | | |
|---|---|---|---|---|
| *HLA-A* | *HLA-B* | *HLA-C* | *HLA-E* | *HLA-F* |
| *HLA-G* | | | | |

| 13-gene MHC class II | | | | |
|---|---|---|---|---|
| *HLA-DMA* | *HLA-DMB* | *HLA-DOA* | *HLA-DOB* | *HLA-DPA1* |
| *HLA-DPB1* | *HLA-DQA1* | *HLA-DQA2* | *HLA-DQB1* | *HLA-DQB2* |
| *HLA-DRA* | *HLA-DRB1* | *HLA-DRB5* | | |

**f**

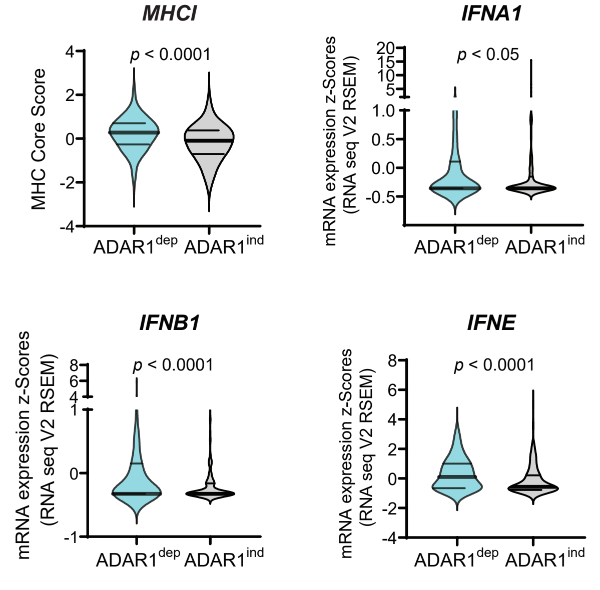


**g**

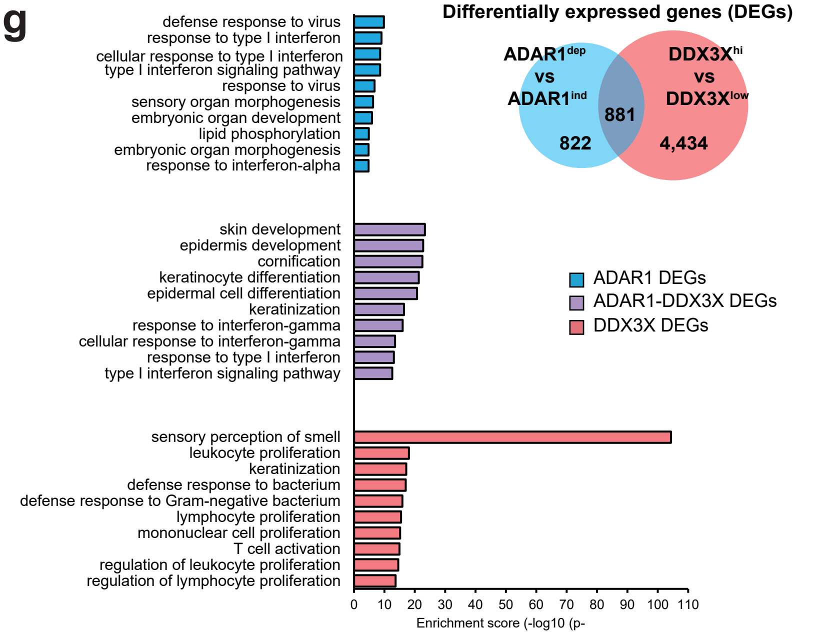

**Extended Fig. 7**

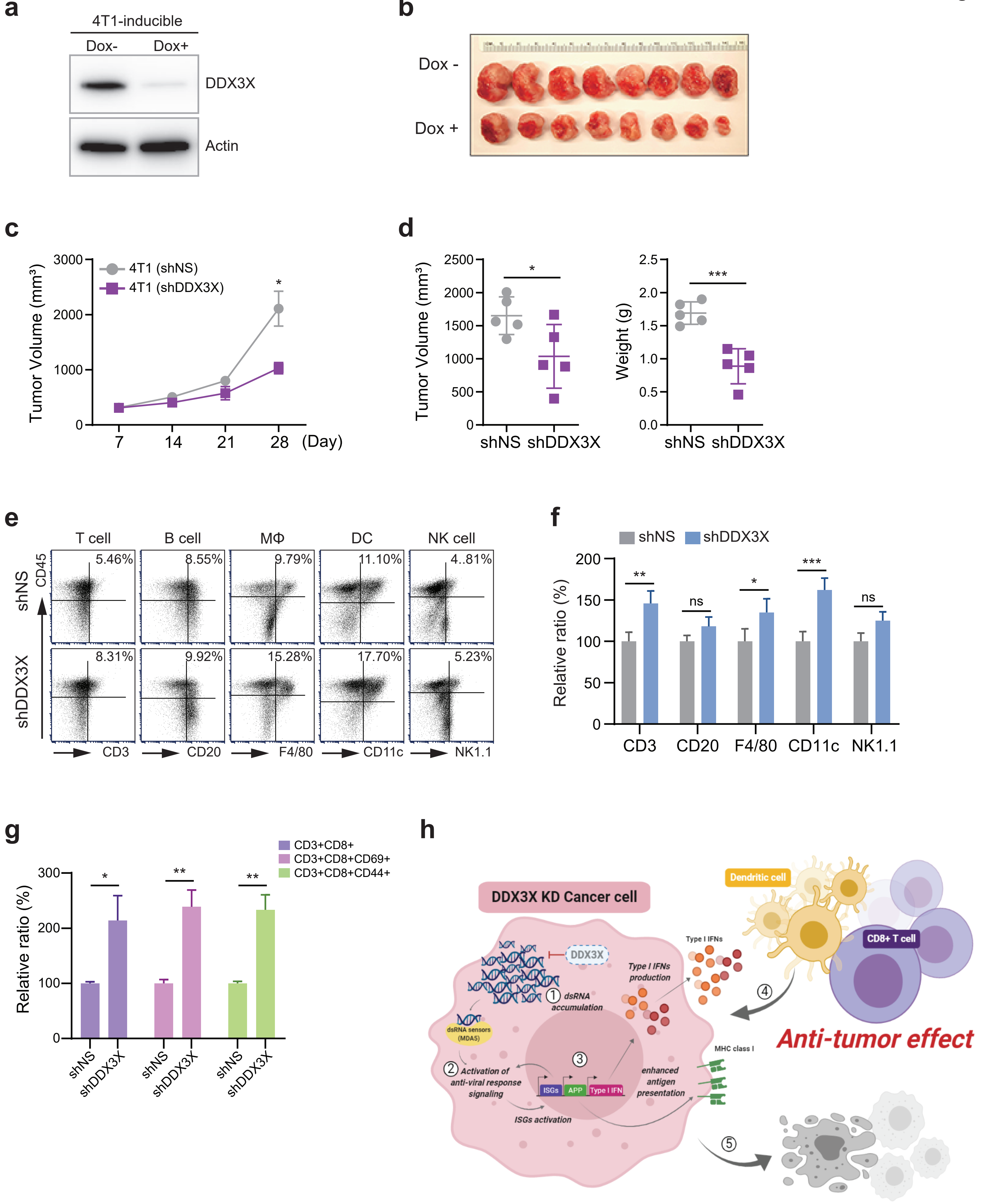

**Extended Data Fig. 1**

**a**, KM-plotter analysis shows the association of DDX3X in breast cancer. High mRNA or protein level of DDX3X is associated with a poor survival outcome in breast cancer. Overall survival, OS; Post-Progression Survival, PPS.

**b**, DDX3X deletion in the MCF7 cells by shRNAs assessed by western blot. A non-specific shRNA (shNS); DDX3X targeting shRNAs (shDDX3X).

**c**, qRT-PCR of gene expression in MCF7 cells treated with siControl (siCtrl) or siDDX3X (left). DDX3X deletion in the MCF7 cells by siRNAs assessed by western blot (right).

**d**, Western blot analysis of DDX1-KD or DDX5-KD MCF7 cells.

**e**, qRT-PCR of mRNA expression of ISGs in DDX3X-control or -KD MDA-MB-453 cells (left). DDX3X deletion in the MDA-MB-453 cells by shRNAs assessed by western blot (right).

Data in C and E are representative of three independent experiments and showed as mean ± SEM. Statistics were calculated using unpaired t-tests. **P* < 0.05; ***P* < 0.01; ****P* < 0.001; ns, not significant.

**Extended Data Fig. 2**

**a**, ELISA of IFN-$\alpha$, -$\beta$, and -$\gamma$ in the culture supernatants from DDX3X-control or -KD A375 cells.

**b**, Expression level of IFN-$\gamma$ and IFN-$\alpha/\beta$ receptors in DDX3X-control or -KD MCF7 or MDA-MB-453 cells.

**c**, Western blot analysis of STAT signaling in MCF7 or MDA-MB-453 cells after 5 ng/ml of IFN-a treatment.

**d**, Protein levels of STAT2 and phosphorylated STAT2 in DDX3X-KD MCF7 or MDA-MB-453 cells.

**e**, Apoptosis analysis of DDX3X-KD MCF7 or MDA-MB-453 cells by flow cytometry. Graphs show the % of early apoptotic cells (positive for annexin-V, negative for sytox-blue; bottom right quadrant) and late apoptotic cells (positive for both annexin-V and sytox-blue; upper right quadrant) for the flow data.

**f**, Colony formation assay in DDX3X-KD MDA-MB-231 or A375 cells shows cell growth inhibition by DDX3X depletion.

**g**, Representative flow histograms and a bar graph of HLA-DR expression on DDX3X-control or -KD MCF7 cells.

Data in A, D, F, and G are representative of three independent experiments. Data are shown as mean ± SEM. Statistics were calculated using unpaired t-tests. **P* < 0.05; ***P* < 0.01; ****P* < 0.001; *****P* < 0.0001; ns, not significant.

**Extended Data Fig. 3**
**a** and **b**, Immunofluorescence analysis (**a**) and dot blotting (**b**) using J2 antibody for endogenous dsRNA in MCF7 cells treated with indicated RNases before analysis.
**c**, Dot blotting of polyI:C using J2 antibody or anti-mouse IgG conjugated HRP ($2^{nd}$ antibody only, upper). J2 dot blotting on RNA extracts from polyI:C transfected A375 cells (lower).
**d**, Schema of DNA methyltransferase inhibitors (5-AzaC) action on ERV production.
**e** and **f**, Increased endogenous dsRNA levels analyzed by dot blot (**e**) and flow cytometry (**f**) after 5-AzaC treatment in each cell. Bar graphs represent relative dot intensity (**e**) and relative MFI (**f**).
**g**, dsRNAs accumulation assessed by dot blot in DDX3X-KD MCF7 cells after 5-azacytidine (5-AzaC) treatment. Treatment with or without 500 nM of 5-AzaC for 3 days and rested for 4 days before assaying.
**h**, Dot blot of RNA extracts from DDX3X-control or -KD HEK293T cells. A Bar graph shows relative dot intensity (right).
**i**, qRT-PCR of ISGs, dsRNA sensing genes (*RIG-I*, *MDA5*), antigen processing and presentation gene (*TAP1*), and *IFNB1* in DDX3X-control or -KD HEK293T cells.
Data are representative of three independent experiments. Data are shown as mean ± SEM. Statistics were calculated using unpaired t-tests. **P* < 0.05; ***P* < 0.01; ****P* < 0.001.

**Extended Data Fig. 4**
**a**, Bar graphs of relative protein band intensity for Fig.4c.
**b**, qRT-PCR of dsRNA sensors, *RIG-I*, *MDA5*, and *OAS1* in DDX3X KD-MCF7 cells.
**c**, Western blot analysis of IRF3 and IRF7 activation after 1ug of polyI:C treatment for 24hr.
**d**, Western blot validation of CRISPR/Cas9-mediated MDA5 KO in DDX3X-control or -KD MCF7 cells.
**e**, Schematic diagram of generating conditional Ddx3x knockout mouse.
**f**, Genotyping of mouse Ddx3x and Ddx3y in tissues and MEF cells.
**g**, Western blot analysis of DDX3X, STAT1, and STAT2 in hemizygous ($Ddx3x^{Y/f}$ male), homozygous ($Ddx3x^{f/f}$ female), and heterozygous ($Ddx3x^{f/+}$ female) MEF cells after Ad-Cre treatment.
**h**, qRT-PCR of *IFNB1* in DDX3X wildtype or $DDX3X^{f/f}$ MEF cells 2 days after Ad-Cre virus treatment.
Data are representative of three independent experiments. Data are shown as mean ± SEM. Statistics were calculated using unpaired t-tests. **P* < 0.05; ***P* < 0.01; ns, not significant.

**Extended Data Fig. 5**

**a**, Validation of cytoplasmic and nucleus isolations from DDX3X-control or -KD MCF7 cells by western blot. (Lamin A/C: nuclear marker, Tubulin: cytoplasmic marker).

**b**, Graphs show the relative RNA concentration and relative dot intensity from Fig. 5a. The relative dot intensity presented compared to DDX3X-control cells (DDX3X-control cells' cytoplasm + nucleus = 100%).

**c**, qRT-PCR analysis for the expression level of mitochondrial genes in the cytoplasm of the DDX3X inducible-KD (Dox+) MCF7 cells.

**d**, Interaction between DDX3X and ADAR1. IP was performed with ADAR1 antibody or control IgG, respectively. Western blot analyzed using anti-DDX3X and anti-ADAR1 antibodies.

**e**, Validation of cytoplasmic and nucleus isolations from MCF7 cells by western blot. (Lamin A/C: nuclear marker, Tubulin: cytoplasmic marker).

**f**, Western blot validation of single or double knockdown of DDX3X and ADAR1 in MCF7 cells.

**g**, qRT-PCR analysis for *DDX3X* and ISGs (*OAS2, IFI44L*) in MCF7 cells with siRNA-mediated knockdown against control, *DDX3X*, and *ADAR1*.

**h**, Schema of A-to I editing assay.

**i**, ADAR1 editing analysis in parent cells.

Data are representative of three independent experiments. Data represented as mean ± SEM. Statistics were calculated using unpaired t-tests. *$P < 0.05$; **$P < 0.01$; ***$P < 0.001$.

**Extended Data Fig. 6**

**a**, Flowchart displaying the steps and database used in this study. RNA-seq and gene dependency data were obtained from Cancer Cell Line Encyclopedia (CCLE) (https://portals.broadinstitute.org/ccle) and DeMap portal (https://depmap.org/portal/), respectively. Immune-related genes were selected based on innateDB (https://www.innatedb.com). RNA-seq data was normalized with the voom method and differential expression determined by limma.

**b**, Cancer type and sex of cancer cells in $DDX3X^{hi}$ and $DDX3X^{low}$ group.

**c**, DDX3X transcript level in each group defined by DDX3X level and sex.

**d**, MHC genes used in calculation of MHC core score.

**e**, Violin plots showing transcript levels of MHCII, IFN, interleukin (IL) genes in $DDX3X^{hi}$ and $DDX3X^{low}$ group.

**f**, Transcript levels of MHC I and type I IFN genes in $ADA1R^{dep}$ versus $ADAR1^{idp}$ group.

**g**, Enrichment of biological processes (GO) terms of total DEGs in ADAR1, ADAR1-DDX3X, and DDX3X group (FDR<0.05) (left). Bars indicate statistical significance shown as −log10 of *p* value. Venn diagram depicting number of DEGs between two groups (right).

**Extended Data Fig. 7**

**a**, Western blot analysis of inducible knockdown of DDX3X in 4T1 cells expressing doxycycline-inducible DDX3X shRNA after incubating 2μg/ml of doxycycline for 72 hours.

**b**, Images of Fig. 6a tumors isolated from the mice at the end point.

**c**, Tumor growth in mice implanted 4T1 cells expressing DDX3X-control (shNS) or DDX3X-knockdown (shDDX3X) shRNAs into mammary fat pad of BALB/cJ.

**d**, Tumor volumes and weights measured after tumor dissection.

**e**, Representative dot plots showing each cell markers expression of CD45+ cells isolated from tumor. CD3+; T cell, CD20+; B cell, F4/80+; Macrophage, CD11c; dendritic cell, NK1.1; NK cell

**f** and **g**, Bar graphs present the frequencies of each cells after flow cytometry analyzed from Extended Data Fig. 7c tumors.

**h**, Illustration of DDX3X depletion mediated anti-tumor effect.

Data are represented as mean ± SD. n=5 mice per group. Unpaired t-tests. $^{*}P < 0.05$; $^{**}P < 0.01$; $^{***}P < 0.001$; ns, not significant.

### Extended Experimental Procedures

#### MEF isolation and conditional DDX3X knockout MEF generation

MEF isolation was performed following to the methods previously described (Durkin et al., 2013; Xu, 2005). In briefly, mouse breeder pairs were set up with one adult male and one to two adult female mice. Female mice were checked copulation plugs in the vagina the following morning. Once plugged female was checked, the mice were removed from breeder cage and keep for 13 days. The pregnant female mouse was euthanized using $CO_2$ and gently pull out embryos encased in uterus. The embryos removed from yolk sac, uterus, and placenta and cut away head, liver and heart. In the new petri dish, the embryo was minced into small pieces. Add 1 ml of trypsin-EDTA into the minced embryo dish and incubate in 37°C incubator for 20 minutes. Add 4 ml of MEF media (DMEM containing high glucose and L-glutamine supplemented with 10% FBS and 1% penicillin/streptomycin) to quench trypsin activity and re-suspended several times to break up tissues. Transfer the cell suspension to 75T flask containing 10 ml of MEF media and placed in 37°C incubator. After cells were attached changed to fresh media and allow cells to grow in 37°C incubator. To remove functional DDX3X, DDX3X wildtype and $DDX3X^{f/f}$ MEFs were treated with Adeno-Cre virus (University of Iowa; Iowa City, IA) multiplicity of infection (MOI) dependently (0-200MOI) for 24 hours and recovered for 2 or 4 days before further analysis.

#### Cytoplasmic and nucleus extractions and analysis

Cytoplasmic and nucleus fractionations were performed using the manufacturer's protocol with slight modification (78833, Thermo Scientific). Cells were harvested with trypsin-EDTA and then centrifuge at 500 x g for 5 minutes. The cell pellet was washed with PBS and removed residual PBS completely. Add appropriate volume of CER I buffer containing protease inhibitor cocktail (87785, Thermo Fisher) to the pellet and suspend the cell pellet carefully less than five times. Cells were incubated for 10 minutes on-ice and then added ice-cold CER II buffer and suspended carefully for 1-2 times. Centrifuged at 16,000 x g for 5 minutes and removed supernatant (cytoplasmic extract) to a clean chilled tube. The pellet was washed with PBS for two times and remove residual PBS completely. The pellet was suspended in NER buffer containing protease inhibitor cocktail and place on-ice for 40 minutes. The supernatant (nuclear extract) was collected after centrifuging at 16,000 x g for 10 minutes. The extracts were used to perform western blot, immunoprecipitation, and RNA isolation for dot blot.

**ELISA**

The supernatant of cells was collected 72 hours after incubation at 37 °C, and the concentration of IFNs were determined using the Human IFN α (41100-1, pbl assay science), Human IFN β (DIFNB0, R&D systems), Human IFN γ (DY285B, R&D systems), Mouse IFN α (42120-1, pbl assay science), Mouse IFN β (DY8234-05, R&D systems), and Mouse IFN γ (DY485-05, R&D systems) ELISA Kit. The ELISA were performed according to the manufacturer's instructions.

**Dual reporter assay**

The single or double knockdown cells of DDX3X and ADAR1 were transfected with the ADAR1 editing reporter plasmids (including positive and negative controls) using ViaFect transfection reagent (E4981, Promega). Reporter activity was measured using Nano-Glo Dual-Luciferase Reporter Assay System (N1610, Promega) according to the manufacturer's instructions.

**Cell colony formation**

Equal numbers of cells were seeded into 6 well plates in triplicate. Cells were fixed with 3.7% formaldehyde in PBS and permeabilized with 100% methanol followed by stained with 1% crystal violet. After staining, cells were washed three times with PBS and dried completely. The cell proliferation was measured by intensity of crystal violet stained colony by Image J (NIH).

**Apoptosis analysis by flow cytometry**

After incubating the cells for 48 hours at 37°C, collect the supernatant containing floating cells and the adherent cells by trypsin EDTA treatment. Cells were stained with Annexin V conjugated with Alexa Fluor 647 (640943, BioLegend) and Sytox blue (425305, BioLegend) in suspend Annexin V binding buffer (422201, BioLegend) for 15 minutes at room temperature in dark. Flow cytometry were performed with a FACSCalibur (BD Biosciences) and data were analyzed using FCS express 6.

**RNA isolation and RNA-Seq Data analysis**

Total RNA was prepared using TRIzol Reagent (15596026, Thermo Fisher) according to the manufacturer's protocols. RNA-seq libraries were prepared using the SOLiD Total RNA-Seq Kit (4452437, Applied Biosystem) and the library quality was checked using Agilent 2100 Bioanalyzer. RNA sequencing was performed on an 5500XL SOLiD Sequencer (Applied Biosystem) according to the manufacturer's protocols. Reads were mapped to the human genome and genic read quantified using LifeScope Genomic Analysis Software and GRCh37 (hg19) genome and transcriptome annotations. Normalization, differential expression analysis, and principle component analysis were performed using R package DEseq2 version 1.45 (Love et al., 2014). Heatmap was constructed using R package pheatmap version 1.0.10. Differentially expressed genes (DEGs) were selected above FDR $< 0.05$ and fold change $> 1.5$. DEGs were used for input into Ingenuity Pathway Analysis (IPA). Gene Set Enrichment Analysis (GSEA) was performed using GSEA software with default setting and associated Molecular Signature Database (MSigDB) as previously described (Subramanian et al., 2005).

**TASA-TD strand-specific PCR**

First strand cDNA synthesis and strand specific PCR for detection of sense and antisense ERV transcripts using TASA-TD methodology was performed according to Henke et al, 2015 (Henke et al., 2015). Primer sequences are shown in Table S1. Specific components from the SuperScriptIII First-Strand Synthesis System for RT-PCR (Life technologies) were used to perform reverse transcription with RNA from MCF7 cells. For the first strand cDNA synthesis reaction, 50 ng of RNA for β-*actin*, 400 ng for *Syncytin-1*, and 500 µg for *Env9-1* were used. 1 uM of a gene specific primer ligated to a TAG-sequence not specific for the human genome (GSP sense/antisense (RT) TAG) was implemented in the reaction. RNA and primers were preheated at 65°C for 5 minutes. For the total reaction: the GSP-TAG, 0.5 mM dNTP, 5 mM $MgCl_2$, 10 mM DTT, 40 U RNaseOUT, 100 U SuperScriptIII RT, and 240 ng Actinomycin D (Sigma) were added with the RNA for a 20 µl reaction. Synthesis was performed at 50°C for 50 minutes and terminated at 85°C for 5 minutes. RT with extremely low intrinsic RNase H activity (for cleavage of RNA from RNA/DNA duplexes) and Actinomycin D was added to prevent second strand cDNA RT resulting in antisense artifacts. After cDNA synthesis, 2 U of recombinant RNase H (life technologies) was added to each reaction and incubated for 20 minutes at 37°C. Afterwards, gene and strand specific PCR was performed. PCR reactions were implemented with the EmeraldAmp GT PCR Master Mix (TAKARA) as described above (PCR analysis). To amplify sense and antisense cDNA, a TAG-primer and GSP sense (PCR) or a TAG-primer and GSP antisense (PCR) were used, respectively. We performed sense and antisense specific PCR using both sense and antisense cDNA of β-*actin* as an internal negative control that was previously demonstrated to have no antisense transcript (Chen et al., 2004). All cDNA products were electrophoresed on 1% agarose gels, and visualized using Gel Red (Biotium). Primer sequence information is available in Supplemental Table 2.

### Extended References


Chen, J., Sun, M., Kent, W.J., Huang, X., Xie, H., Wang, W., Zhou, G., Shi, R.Z., and Rowley, J.D. (2004). Over 20% of human transcripts might form sense-antisense pairs. Nucleic Acids Res *32*, 4812-4820.

Durkin, M.E., Qian, X., Popescu, N.C., and Lowy, D.R. (2013). Isolation of Mouse Embryo Fibroblasts. Bio Protoc *3*.

Henke, C., Strissel, P.L., Schubert, M.T., Mitchell, M., Stolt, C.C., Faschingbauer, F., Beckmann, M.W., and Strick, R. (2015). Selective expression of sense and antisense transcripts of the sushi-ichi-related retrotransposon--derived family during mouse placentogenesis. Retrovirology *12*, 9.

Love, M.I., Huber, W., and Anders, S. (2014). Moderated estimation of fold change and dispersion for RNA-seq data with DESeq2. Genome Biol *15*, 550.

Subramanian, A., Tamayo, P., Mootha, V.K., Mukherjee, S., Ebert, B.L., Gillette, M.A., Paulovich, A., Pomeroy, S.L., Golub, T.R., Lander, E.S.*, et al.* (2005). Gene set enrichment analysis: a knowledge-based approach for interpreting genome-wide expression profiles. Proc Natl Acad Sci U S A *102*, 15545-15550.

Xu, J. (2005). Preparation, culture, and immortalization of mouse embryonic fibroblasts. Curr Protoc Mol Biol *Chapter 28*, Unit 28 21.
